# Activating the CXCR3/CXCL10 pathway overrides tumor immune suppression by enhancing immune trafficking and effector cell priming in head and neck squamous cell carcinoma

**DOI:** 10.1101/2025.04.24.650529

**Authors:** Cheyanne K. Shinn, Robert Saddawi-Konefka, Catherina L. Salanga, Shiruyeh Schokrpur, J. Silvio Gutkind, Tracy M. Handel

## Abstract

The immune-suppressive nature of the tumor microenvironment (TME) has limited the impact of immune checkpoint blockade in many cancers, often by restricting the infiltration and activation of anti-tumoral CD8+ T, CD4+ T, and NK cells. Here, we utilized murine models of head and neck squamous cell carcinoma and demonstrated that intratumoral (IT) delivery of CXCL10 drives tumor elimination and inhibits recurrence. CD8+ T cells recruited to tumors display enhanced activation, increased tumor antigen specificity, and decreased markers of T cell exhaustion, indicating that CXCL10 not only directs cell migration, but also enhances T cell effector functions. Despite delivery of CXCL10 into tumors, CD8+ and CD4+ T cells also show enhanced presence and proliferation in tumor-draining lymph nodes (TdLNs), consistent with antigen presentation and trafficking of these cells between tumors and TdLNs. CXCL10 also stunts angiogenesis and lymphangiogenesis within the TME, which likely contributes to its antitumoral effects. Finally, enhanced tumor clearance was observed by combining IT CXCL10 and anti-PD-1. Together, these findings provide the rationale for the clinical evaluation of CXCL10 as a strategy to enhance the efficacy of immunotherapy.

**Graphical Abstract:** 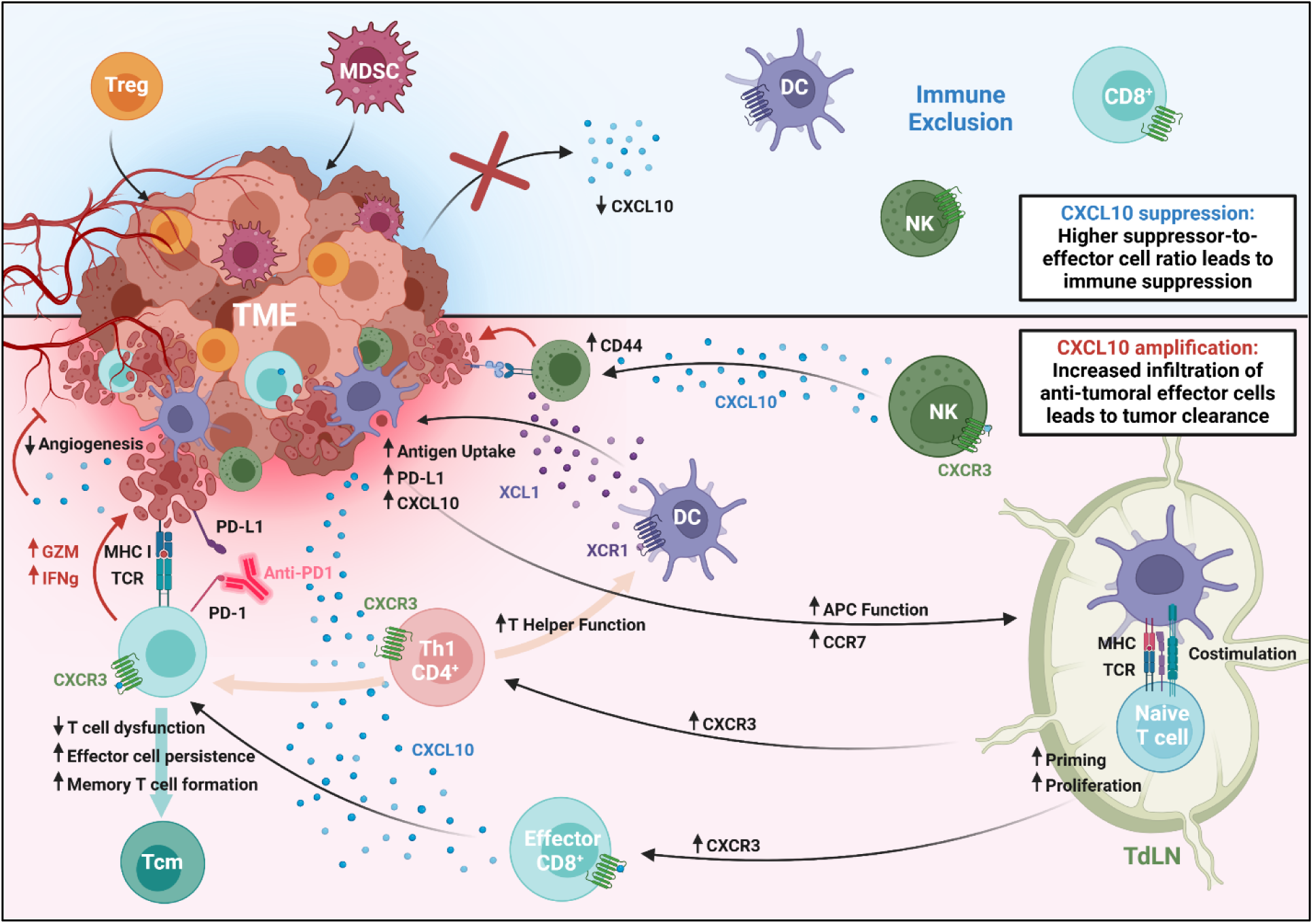

**SUMMARY:** CXCL10 suppresses tumor growth and promotes immune memory by recruiting T and NK cells into the tumor microenvironment, promoting tumor-specific antigen recognition and effector functions, slowing T cell exhaustion, and inhibiting angiogenesis. CXCL10 directly and indirectly mobilizes an immune cell network that together supports an anti-tumoral microenvironment.

## INTRODUCTION

Immunotherapy has revolutionized cancer treatment by leveraging the host immune system to target and eliminate tumors, offering new therapeutic approaches for patients with previously untreatable cancers. In particular, immune checkpoint blockade agents (ICBs) such as pembrolizumab and nivolumab have exhibited success in treating patients with recurrent or metastatic disease in several human cancers (*1*). These agents work by disrupting PD-1/PD-L1 signaling cascades that block the activation and proliferation of cytotoxic T cells, inhibit cytokine production, and accelerate CD8+ T cell exhaustion (*2–4*). However, despite promising results in triggering anti-tumor immune responses, favorable clinical outcomes remain limited and vary drastically across different cancers, with objective response rates of 33–40% in melanoma (*5, 6*), 15–20% in non-small cell lung cancer (*7*), 22–25% in renal cell carcinoma (*8*), and 10-20% in head and neck squamous cell carcinoma (*9–11*). The failure of these ICBs, as well as standard treatments, is due in part to the ability of resistant tumors to limit the infiltration and activation of cytotoxic CD8+ T and NK cells (*12–14*). CD8+ T and NK release apoptosis-inducing cytotoxins like perforin and granzyme, as well as pro-inflammatory cytokines such as interferon gamma (IFN-γ) and tumor necrosis factor alpha (TNF-α) to eliminate cancer (*15*), and are essential for an effective response to immunotherapy. Accordingly, the abundance of cytotoxic cells within tumor tissue following various treatment strategies is associated with a positive prognosis in several cancers (*16–18*). However, exclusion of these cells from tumors can result from physical barriers like dense extracellular matrix deposition (*19*), immunosuppressive signaling from tumor and stromal cells (e.g., TGF-β and VEGF-mediated immune evasion) (*20*), and dysfunctional vasculature that impairs effective immune cell infiltration (*21*). Further, the mere infiltration of cytotoxic cells into the tumor microenvironment (TME) does not guarantee positive outcomes. Immunosuppressive cytokines and chronic antigen stimulation can lead to the dysfunction and exhaustion of these cells, limiting their ability to control tumor growth (*22, 23*). Additionally, the exclusion of dendritic cells (DCs) also contributes to immune suppression by stunting the generation of T cells that are primed to tumor-specific antigens (TSAs) (*24, 25*). Thus, to broaden the success of immunotherapy, approaches are needed that override the obstruction of key anti-tumoral cells and promote the generation and maintenance of T cells that are reactive to TSAs.

The chemokine receptor CXCR3 is a key regulator of cytotoxic immune cell responses within solid tumors (*26, 27*). It responds to three interferon inducible ligands (CXCL9, 10, and 11) and controls the migration, spatial distribution, and function of activated T cell and NK cells (*28, 29*). CXCL9 and 10 have been shown to be particularly important for the anti-tumoral functions of CD8^+^ T cells, Th1 CD4^+^ T cells, and NK cells (*30–32*) and reduced expression of either ligand contributes to immune exclusion within the TME (*26, 33, 34*). Conversely, high expression of CXCL9, CXCL10, and their receptor, CXCR3, is associated with improved responses to ICB treatment in several human cancers and murine cancer models (*35–37*). CXCL10 has been shown to impact the polarization of CD4^+^ T cells into anti-tumoral Th1 subtypes, reflecting its ability to influence T cell differentiation in addition to guiding migration (*38*). It has also been shown to play a key role in facilitating interactions between CD8+ T cells and antigen-presenting DCs, fostering a pro-inflammatory niche that counters immunosuppressive factors and leads to effective T cell activation, expansion, and tumor elimination (*39, 40*). These findings suggest that by increasing intratumoral CXCL10 levels, it may be possible to override immunosuppressive mechanisms and reprogram the TME to a more antitumoral state responsive to ICB immunotherapy.

To explore this hypothesis, we focused on head and neck squamous cell carcinoma (HNSCC), a highly prevalent and aggressive cancer with limited immunotherapeutic success. HNSCC is a significant global health concern, ranking seventh in cancer incidence and accounting for more than 700,000 new cases diagnosed annually (*41*). Approximately 75% of HNSCCs are associated with tobacco and alcohol use (*42, 43*), though human papilloma virus (HPV) infection is rising as the major cause of oropharyngeal squamous cell carcinoma (*44, 45*). Patients with HPV-negative, tobacco-related HNSCC undergo aggressive treatments involving surgery, radiation, and chemotherapy, yet the 5-year survival rate remains at 40-50% (*46*), and only 10-20% respond to PD-1/PD-L1 blockade (*9, 10*), underscoring the need for new therapeutic strategies. The abundance of cytotoxic cells within tumor tissue following various treatment strategies is associated with a positive prognosis in HNSCC (*17, 18*). Conversely, the loss of chromosome segment 9p21/9p24 in HPV-negative patients results in immunologically cold ICB-resistant HNSCC tumors, which correlates with a profound decrease in CXCL9 and CXCL10 expression and markedly reduced patient outcomes (*47, 48*).

Here, we leverage the 4NQO-induced Murine Oral Squamous Cell (4MOSC1) carcinoma (*49*) and Murine Oral Carcinoma (MOC1) (*50*) mouse models of HNSCC to investigate the efficacy of IT delivery of CXCL10. We show that CXCL10 alone suppresses tumor growth and recurrence by enhancing the infiltration, activation, and effector function of CD8+ T cells, CD4+ T cells, and NK cells. CD8+ T cells acquired the ability to recognize tumor-specific antigens, which correlated with their migration to and proliferation in tumor-draining lymph nodes (tdLNs), despite the administration of CXCL10 into tumors. This, along with structural changes such as reduced angiogenesis and lymphangiogenesis, suggests that CXCL10 directly and indirectly mobilizes a network of immune cells that support a multimodal program of anti-tumoral activities, extending beyond the TME. Combining IT CXCL10 with anti-PD-1 therapy effectively doubled the number of mice that completely cleared tumors and showed durable responses. These findings provide the rationale for the clinical evaluation of IT CXCL10 as a strategy to overcome tumor-regulated immunosuppression, restoring sensitivity to ICB immunotherapies, and improving outcomes for HNSCC patients.

## RESULTS

### High CXCR3 and CXCL10 expression correlates with early tumor stage, increased immune infiltration, inflammatory gene signatures, and improved overall outcomes in patients with HNSCC

To determine the impact of CXCR3 and CXCL10 on overall outcomes and inflammatory status in HNSCC patients, we first interrogated the Cancer Genome Atlas (TGCA, PanCancer Atlas) with cBioPortal (http://cbioportal.org) (*51, 52*). Tumors with high *CXCR3* and *CXCL10* mRNA expression correlated with increased overall survival in patients with HNSCC as characterized by hazard ratios (HR) less than 0.5 and 0.73, respectively (**Fig. 1A-B**). These results are consistent with prior findings demonstrating that tumors with elevated co-expression of *CXCL10* and signal transducer and activator of transcription 2 (*STAT2*) are associated with better outcomes of oral cancer patients (*53, 54*). Furthermore, high *CXCR3* and *CXCL10* mRNA expression in HNSCC tumors show a positive correlation with CD8 antigen (*CD8A*) (**Fig. 1C-D**) indicating an elevated presence of CD8^+^ cells. Bulk mRNA expression data was also utilized to gauge changes in the inflammatory signatures of the CXCL10^high^ vs CXCL10^low^ cohorts. These data showed that the expression levels of both interferon gamma (*IFNG*), a pleiotropic cytokine involved in multiple adaptive and innate inflammatory responses, and granzyme B (*GZMB*), a serine protease indicative of cytotoxic activity, were significantly higher in the CXCL10^high^ tumors (**Fig. 1E**). IFN-γ is considered one of the main inducers of PD-L1 expression in cancer cells, rationalizing associations between *IFNG* expression or IFN-γ inducible gene signatures and positive clinical responses to PD-1/PD-L1 blockade in HNSCC patients (*35, 55*) as well as melanoma (*56*) and bladder carcinomas (*57*). Additional differentially expressed genes associated with interferon-mediated inflammatory responses were revealed in the CXCL10^high^ vs CXCL10^low^ cohorts, including interferon-induced CXCR3 chemokines (*CXCL9* and *CXCL11*), *STAT1/2* (*58*), interferon regulatory factor 5 (*IRF5*) (*59*), and interferon-induced guanylate-binding proteins (*GBP1/4/5*) (*60*) (**Fig. 1F, Fig. S1**). The differential expression of transporter associated with antigen processing 1 (*TAP1*) (**Fig. S1**), a protein involved in the processing and presentation of major histocompatibility complex class I (MHC1) restricted antigens for recognition by CD8+ T cells (*61*), provides additional evidence of high *CXCL10* expression correlating with increased cytotoxic activity. HNSCC lesions are often classified as early-stage (stage I–II) and late-stage (stage III– IV), based on tumor size and local invasion (*62*). A comparison of stage distribution amongst HNSCC tumors in CXCL10^high^ vs CXCL10^low^ cohorts showed that 62.85% of tumors in the CXCL10^low^ cohort were graded in late-stage III–IV, whereas 44.87% of CXCL10^high^ tumors were enriched in early-stage disease (**Fig. 1G**). These findings reinforce the role of CXCR3/CXCL10 signaling in regulating tumor-immune suppression and disease progression.

**Figure 1.**
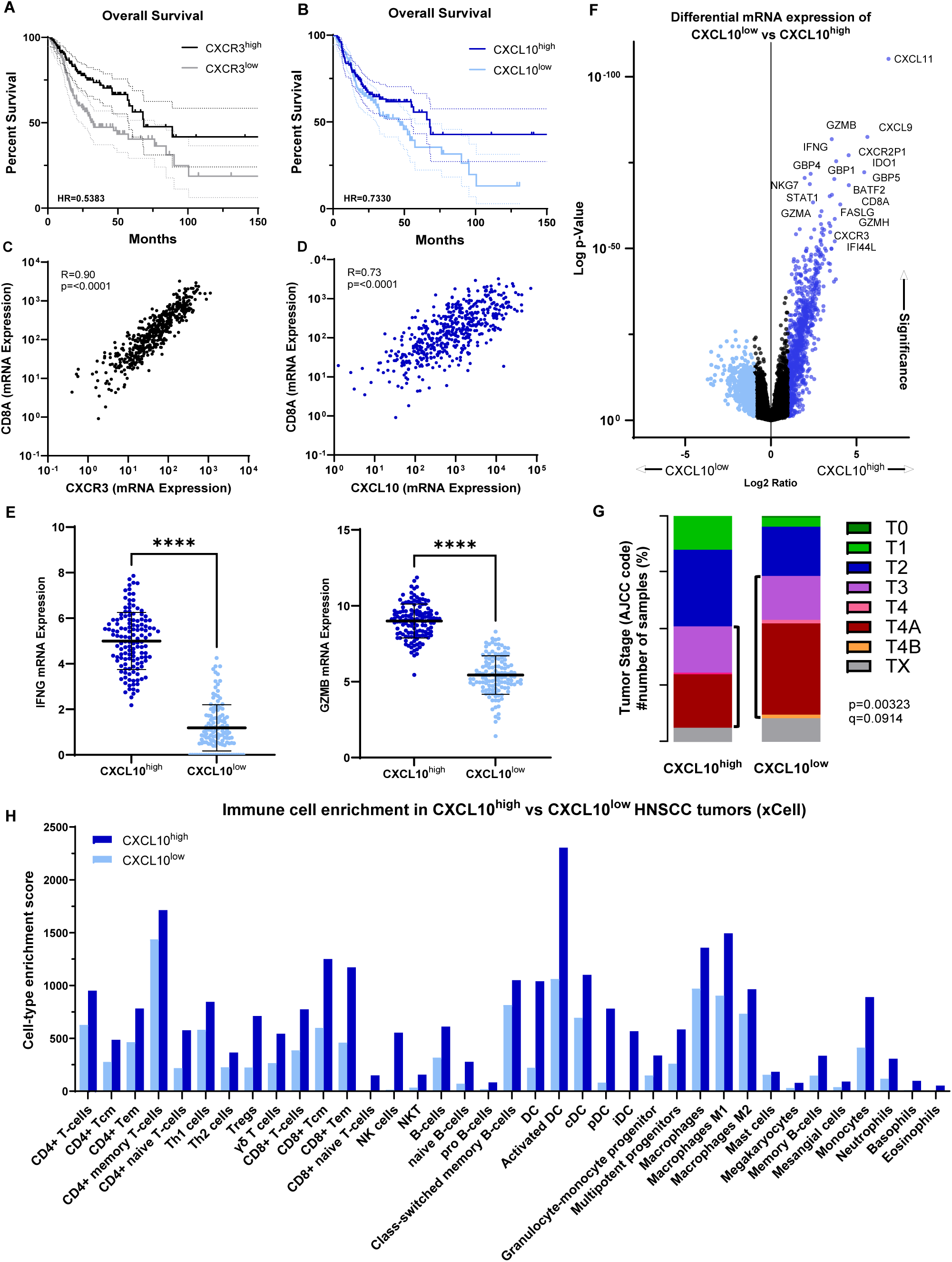
High *Cxcr3* and *Cxcl10* expression correlates with increased immune infiltration into tumors and improved outcomes of patients with HNSCC. (**A-G**) Transcriptomic data obtained and analyzed via cBioPortal (PanCancer Atlas, HNSCC cohort). Expression levels are based on whole tumor mRNA quantification. CXCR3^high^/CXCL10^high^ cohorts (*n* = 128 and 132) and CXCL10^low^/CXCL10^low^ cohort (*n* = 131 and 126) represent the upper 25% and lower 25% threshold for *Cxcr3* or *Cxcl10* mRNA expression, respectively. (**A**) Overall survival of CXCR3^high^ vs CXCR3^low^ cohorts to month 150, with hazard ratio of 0.5383 and p-value of 0.0016 (via Mantel-Cox and Mantel-Haenszel tests). (**B**) Overall survival of CXCL10^high^ vs CXCL10^low^ cohorts to month 150, with hazard ratio of 0.7330 and p-value of 0.1115 (via Mantel-Cox and Mantel-Haenszel tests). (**C-D**) Correlation of *Cxcr3* (**C**) and *Cxcl10* (**D**) with *Cd8a* mRNA expression (Spearman’s rank correlation). (**E**) Relative *Ifng* and *Gzmb* mRNA expression of CXCL10^high^ vs CXCL10^low^ cohorts. (**F**) Differential expression of genes between CXCL10^high^ vs CXCL10^low^ cohorts using Wilcoxon Rank Sum test. Log_2_ fold-change and P value (adjusted) cut-offs are 1 and 0.05, respectively. (**G**) AJCC tumor stage distribution of patients in CXCR3^high^ and CXCR3^low^ cohorts. (**H**) Immune enrichment scores for indicated immune cell subtype in CXCR3^high^ and CXCR3^low^ cohorts.

To explore the association between immune infiltration profiles and the average expression of the CXCL10^high^ vs CXCL10^low^ cohorts, cell enrichment profiles were also derived from TCGA gene expression data using the xCELL platform (*63*) (**Fig. 1H**). Elevated expression of *CXCL10* in HNSCC patients is associated with a broad increase in immune infiltration, with over 2-fold increases in CD8^+^ T cell subtypes, particularly antigen-experienced central (T_CM_) and effector (T_EM_) memory CD8+ T cells. These cells are capable of rapid proliferation and differentiation into effector T cells, which enable a faster and more robust immune response upon re-exposure to tumor antigen (*64*). CXCL10 expression also showed a very strong correlation with NK cell (>40-fold) and Natural Killer T cell (NKT) signatures (>4-fold) (**Fig. 1H**); NKT cells are a subset of CD1d- restricted T cells that have characteristics of both NK and T cells and act as an interface between the innate and adaptive immune system (*65*). DCs not only serve as the primary producers of CXCL10, but are also required for the presentation of tumor-derived antigens to CD4^+^ and CD8^+^ T cells and their priming and activation (*25, 66*). Notably, the data showed a correlation between high CXCL10 expression and broad DC gene signatures, including activated DCs (>2-fold), plasmacytoid DCs (pDCs, which express CXCR3) (>9-fold), and immature DCs (iDCs) (>500-fold) (**Fig. 1H**). These findings further indicate that increased CXCL10 expression has a strong association with anti-tumor immune signatures, particularly favoring the increased infiltration of cytotoxic effector cells and antigen-presenting DCs.

### Intratumoral CXCL10 suppresses tumor growth and recurrence in syngeneic mouse models of HNSCC

Given the correlation between positive outcomes in HNSCC patients and high expression of CXCL10, we hypothesized that elevating tumor-localized CXCL10 levels by IT injection would promote tumor rejection in mouse models of HNSCC. Localized IT delivery rather than systemic administration is required for chemokine agonists in order to direct the migration of cells into the tumor. For initial studies, we employed the 4MOSC1 model as tumors from this model share 98.9% similarity to human tobacco-associated HNSCC based on exome sequencing and mutational signatures, as well as similar immune infiltrates, and limited clinical responses to immunotherapy (*49*). 4MOSC1 tumors can be established orthotopically in the tongue or buccal space of mice, both of which are amenable to IT injection of CXCL10 (*67*). Thus, the 4MOSC1 mouse model provides an excellent platform for studies that seek to reprogram the cellular and inflammatory status of the TME with exogenous ligand addition. As these experiments required large quantities of highly purified chemokine, we modified a previously described *E. coli* expression/purification system (*68*) to optimize the efficient production of murine CXCL10 with high yield (∼13.5 mg/L) and purity (**Fig. S2A-E**).

As described in the Methods section, 4MOSC1 tumors were established by injecting one million 4MOSC1 cells into the buccal space of 6-8 week-old C57BL/6 mice. CXCL10 (10 µg) was administered intratumorally on days 3 and 6 post-tumor engraftment (**Fig. 2A**). This treatment schedule was informed by our prior observations that highlight a critical timeframe for development of adaptive immune responses in the 4MOSC1 model: specifically, early lymphablation (days 3 and 6 post-tumor engraftment) eliminated responses to ICB and worsened overall survival by limiting antigen-specific CD8-driven immunity whereas late lymphablation (beyond day 11) did not impact the effectiveness of ICB (*67*). As shown in **Fig. 2B and 2C**, all mice treated with CXCL10 exhibited a significant reduction in tumor growth, with 30% (**Fig. 2C**, **Fig. S2F**; 24.4% of 4MOSC1 mice on average over all experiments) showing complete resolution and remaining tumor-free six weeks after clearance. Subsequent rechallenge with 4MOSC1 cells six weeks after the initial tumor clearance did not result in tumor formation, indicating the formation of durable immunological memory (**Fig. 2D**).

**Figure 2.**
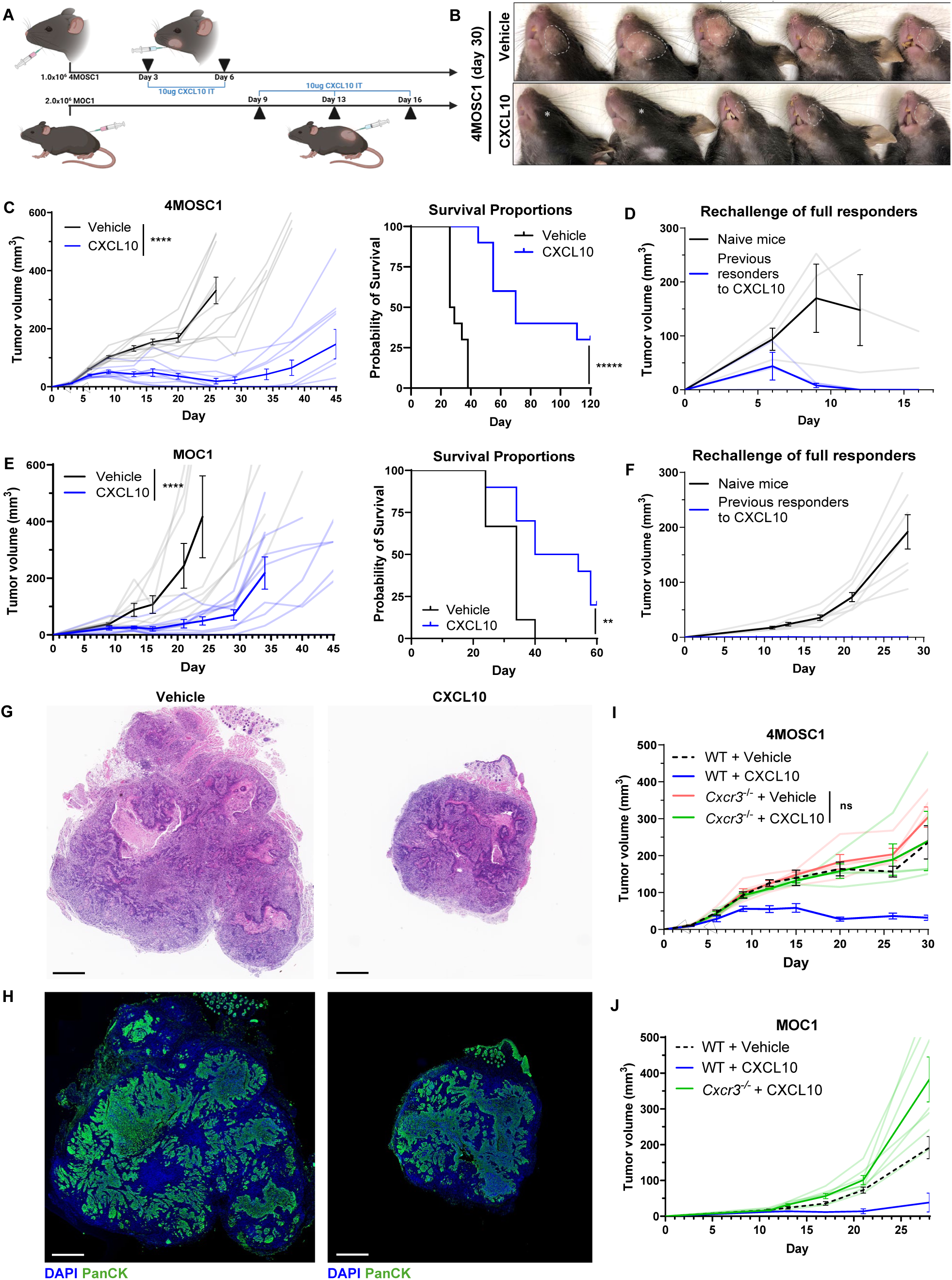
IT delivery of CXCL10 suppresses tumor growth and reoccurrence in syngeneic models of HNSCC. (**A**) Schematic detailing experimental setup and dosing schedule for 4MOSC1 and MOC1 models. 1.0 x 10^6^ 4MOSC1 or 2.0 x 10^6^ MOC1 cells were implanted into the buccal space or flank, respectively, of C57BL/6 mice on day 0. Mice received either 10 µg CXCL10 in 50 µL of PBS (per dose) or 50 µL of PBS (vehicle) on the days indicated (▴). (**B**) Representative image of 4MOSC1 buccal tumors in vehicle vs CXCL10-treated mice at day 30. Tumor boundaries are traced with a dotted line, while an asterisk (*) indicates complete tumor clearance. 4MOSC1 (**C**) and MOC1 (**E**) tumor kinetics of vehicle vs CXCL10-treated mice (*n* = 9-10 per group). Survival proportions are shown on the right. 4MOSC1 (**D**) and MOC1 (**F**) tumor kinetics of tumor naïve age-matched mice or previous complete responders to CXCL10 treatment after rechallenge with 1.0 x 10^6^ 4MOSC1 or 2.0 x 10^6^ MOC1 cells 6 weeks post-tumor clearance. (**G-H**) Representative H&E (**G**) and IF (**H**) staining of 4MOSC1 tumors from vehicle vs CXCL10- treated mice at day 10. Scale bar, 800 μm. (**I-J**) Tumor kinetics of WT or *Cxcr3*^-/-^ mice receiving either vehicle or CXCL10 treatment. Differences between the tumor kinetics amongst experimental groups were analyzed by linear regression. Survival analysis was performed using the Kaplan–Meier method and log-rank tests. All data represent averages ± SEM, except where indicated.

To explore the generalizability of these findings in other syngeneic models of murine HNSCC, we also used the Murine Oral Carcinoma (MOC1) model, which like the 4MOSC1 model, exhibits a partial response to PD-1/PD-L1 blockade that is comparable to patient outcomes (*69*). Following detection of MOC1 tumors in the flank at day 7 post-tumor engraftment, CXCL10 (10 µg) was administered on days 9, 13, and 16 (**Fig. 2A**). Consistent with the 4MOSC1 model, a significant suppression of tumor growth was observed in all CXCL10-treated MOC1 tumors, with 22.2% (**Fig. 2E**, **Fig. S2F**; 21.9% of 4MOSC1 mice on average over all experiments) of the mice demonstrating a complete response and quickly clearing subsequent tumors upon rechallenge (**Fig. 2F**). While smaller in overall size, 4MOSC1 tumors from CXCL10-treated mice exhibit typical HNSCC histology, as indicated by hematoxylin and eosin (H&E) and pan cytokeratin (panCK) (**Fig. 2G-H**). Finally, to confirm the specificity of targeting CXCR3-expressing cells, we performed experiments utilizing CXCR3-deficient (CXCR3^-/-^) mice in both the 4MOSC1 and MOC1 models. Mice lacking CXCR3 failed to eliminate tumors following treatment with CXCL10 (**Fig. 2I-J**), demonstrating a direct role of CXCR3 activation in the anti-tumoral responses to CXCL10 treatment.

### CXCL10 remodels the tumor immune microenvironment by enhancing the infiltration of CD8+ T, CD4+ T, and NK cells

Following activation, naïve T cells upregulate CXCR3 expression, which remains persistent on effector CD8^+^ T cell as well as type-1 helper (Th1) CD4^+^ T cells (*70, 71*) that support CD8+ T cell functions (*72, 73*). CXCR3 is also highly expressed on innate NK cells and NKT cells, and is crucial for the localization of these first-line defenders to sites of inflammation and tumors (*33, 74*). Thus, we hypothesized that CXCL10 treatment would promote the infiltration of these cell types within tumors. To explore changes in the immune status of the TME induced by IT CXCL10 administration in 4MOSC1 tumors, we employed the nCounter PanCancer Mouse Immune Profiling gene expression platform (NanoString Technologies), which surveys 770 immune-related gene signatures, thereby enabling broad characterization of immune cell populations and inflammatory status. As shown in **Fig. 3A**, CXCL10 treatment resulted in elevated T and cytotoxic cell mRNA signatures, including signatures specific to CD8^+^ T cells and NK cells compared to vehicle treatment. To confirm whether the changes in the TME are dependent on these immune subtypes, we treated mice bearing 4MOSC1 tumors with CXCL10 and either CD8-depleting, NK1.1-depleting, or CD4-depleting antibodies. While a partial loss of CXCL10 response was observed with depletion of both CD4^+^ and NK cells, CD8 depletion resulted in the rapid and robust growth of 4MOSC1 tumors despite CXCL10 treatment, highlighting the critical role that CD8^+^ T cells play in the tumor immune response to CXCL10 (**Fig. 3B**). Subsequent analysis by flow cytometry further validated these findings: tumors from CXCL10-treated mice exhibited increased total CD8^+^ T cell, CD4^+^ T cell, and NK cell numbers starting as early as day 7, which persisted through day 10, compared to vehicle control (**Fig. 3C-D**).

**Figure 3.**
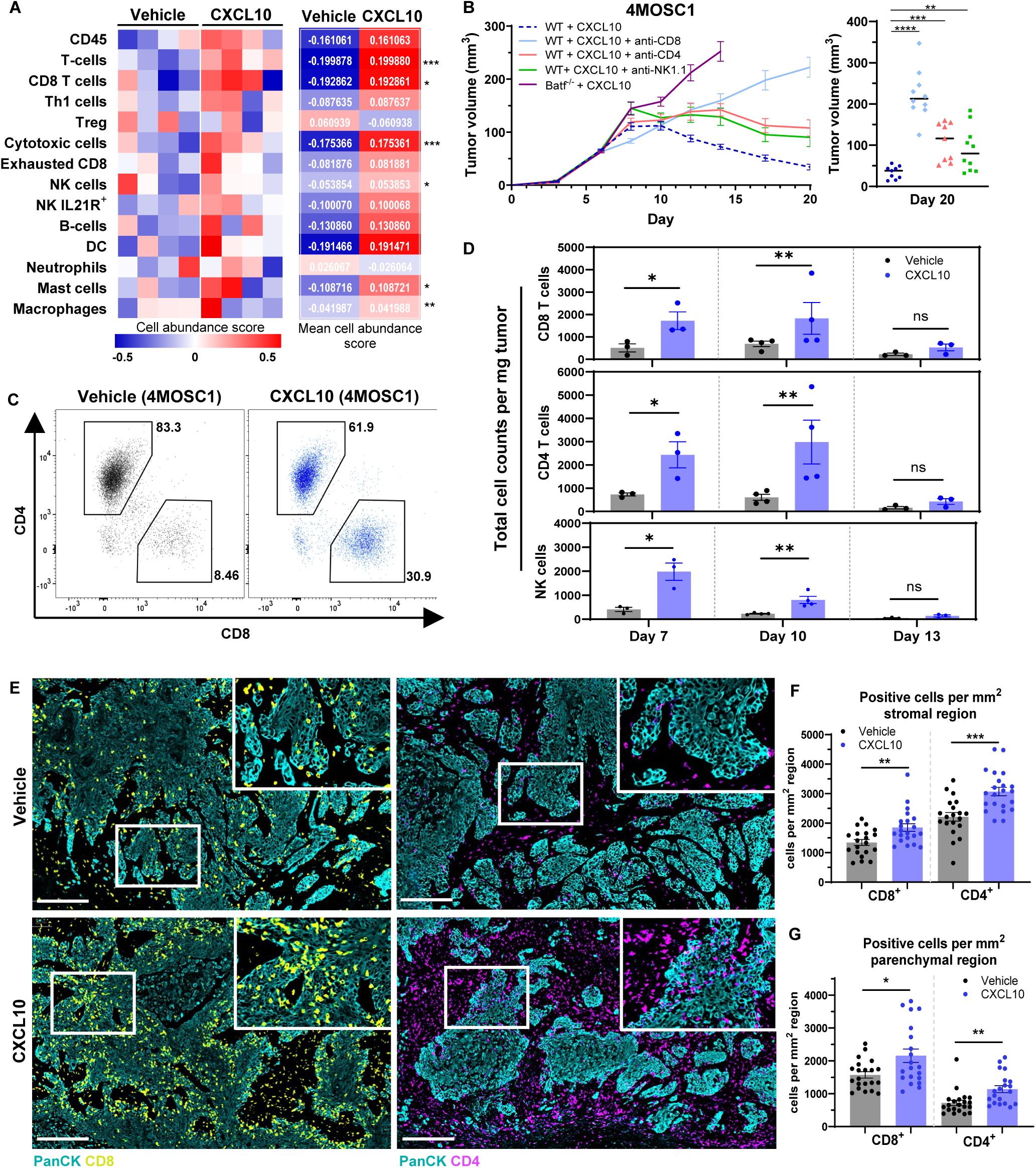
CXCL10 remodels the tumor immune microenvironment by enhancing infiltration of CD8+ T cells, CD4+ T cells, and NK cells. (**A**) 1.0 x 10^6^ 4MOSC1 cells were implanted into the buccal space of C57BL/6 mice on day 0. Tumors were treated on days 3, 6, and 9 with either vehicle or 10 µg of CXCL10 IT (per dose). mRNA from each tumor was isolated and comprehensive immune profiling was conducted using the NanoString nCounter PanCancer Mouse Immune Profiling gene expression platform. Rosalind software was used to analyze genes associated with the listed immune cells and each replicate assigned a cell abundance score (n = 4 mice per group). (**B**) Growth kinetics of 4MOSC1 tumors treated with either vehicle, CXCL10 alone, CXCL10 + anti-CD8, CXCL10 + anti-CD4, or CXCL10 + anti-NK1.1 antibody. Volumes of individual tumors in mm^3^ at day 20 are shown on the right. (**C**) Representative flow cytometry plots showing relative CD4^+^ and CD8^+^ cells amongst total CD3^+^ cells. (**D**) Absolute number of live CD45^+^CD3^+^CD8^+^ T cells, CD45^+^CD3^+^CD4^+^ T cells, and CD45^+^NK1.1+ cells infiltrating vehicle or CXCL10-treated 4MOSC1 tumors at days 7, 10, and 13. Shown is the average number of cells infiltrating per mm^3^ of tumor (*n* = 3 mice per group; multiple unpaired Student’s *t* test; data are represented as mean ± SEM). (**E**) Representative immunostaining of CD8+ and CD4+ cells within the stroma and tumor parenchyma (PanCK) of vehicle and CXCL10-treated tumors at day 10. The scale bar represents 200 μm. (**F-G**) Quantification of total CD8+ (yellow) (**F**) and CD4+ (magenta) (**G**) cells within the tumor stroma and tumor parenchyma (panCK, cyan) (*n* = 4 mice per group; 5-6 regions of interest (ROIs) screened per tumor; ROI designation of stroma vs tumor parenchyma designated based on PanCK expression).

The spatial distributions of tumor infiltrating T cells can be a determinant of overall outcomes and are categorized into three phenotypic categories: immune-desert (lack of leukocyte infiltration), immune-excluded (stromal infiltration without leukocytes entering the tumor parenchyma), and immune-inflamed (leukocyte infiltration beyond the tumor parenchyma) (*75, 76*). In general, immune-inflamed tumors are associated with a more favorable overall prognosis and respond more positively to chemotherapy and ICB (*77*). Thus, we also investigated the abundance and spatial distribution of infiltrating immune cells within tumors via immunofluorescence (IF) staining. At day 10, tumors from CXCL10-treated mice showed substantial increases in infiltrating CD8^+^ and CD4^+^ T cells within the tumor parenchyma and surrounding stroma, suggesting an immune-inflamed tumor phenotype, while tumors from vehicle-treated mice presented a more immune-excluded phenotype (**Fig. 3E-G**). In contrast to CD8^+^ and CD4^+^ T cell infiltration and distribution, most infiltrating NK cells were in the stroma of tumors from CXCL10-treated mice, similar to the vehicle control mice (**Fig. S3A-B**). However, tumors from CXCL10-treated mice also displayed a larger proportion of CD44^+^ NK cells by flow cytometry at day 7, which persisted through day 13 compared to vehicle control (**Fig. S3C**). As increased CD44 expression on NK cells has been attributed to increased cytotoxic activity and IFN-γ production (*78, 79*), these data suggest that CXCL10 treatment induced these cells to acquire activation markers consistent with increased anti-tumoral functions.

### CXCL10 promotes the formation of effector and memory phenotypes in T cells

In addition to the well-established involvement of the CXCR3-ligand system in recruiting cytotoxic effector cells to the TME, there is emerging evidence that the ligands also elicit phenotypic changes in CXCR3-expressing immune cells. For example, CXCL10 has been shown to induce CD4+ T cell polarization towards immune-supporting Th1 and Th17 lineages (*80, 81*), and to promote terminal differentiation of effector CD8+ T cells (*82*), as well as T cell interactions with DCs (*39*). We therefore interrogated phenotypic changes of tumor-infiltrating immune cells in both CXCL10- and vehicle-treated 4MOSC1 tumors via bulk mRNA profiling (**Fig. 4, Fig. S4A**).

**Figure 4.**
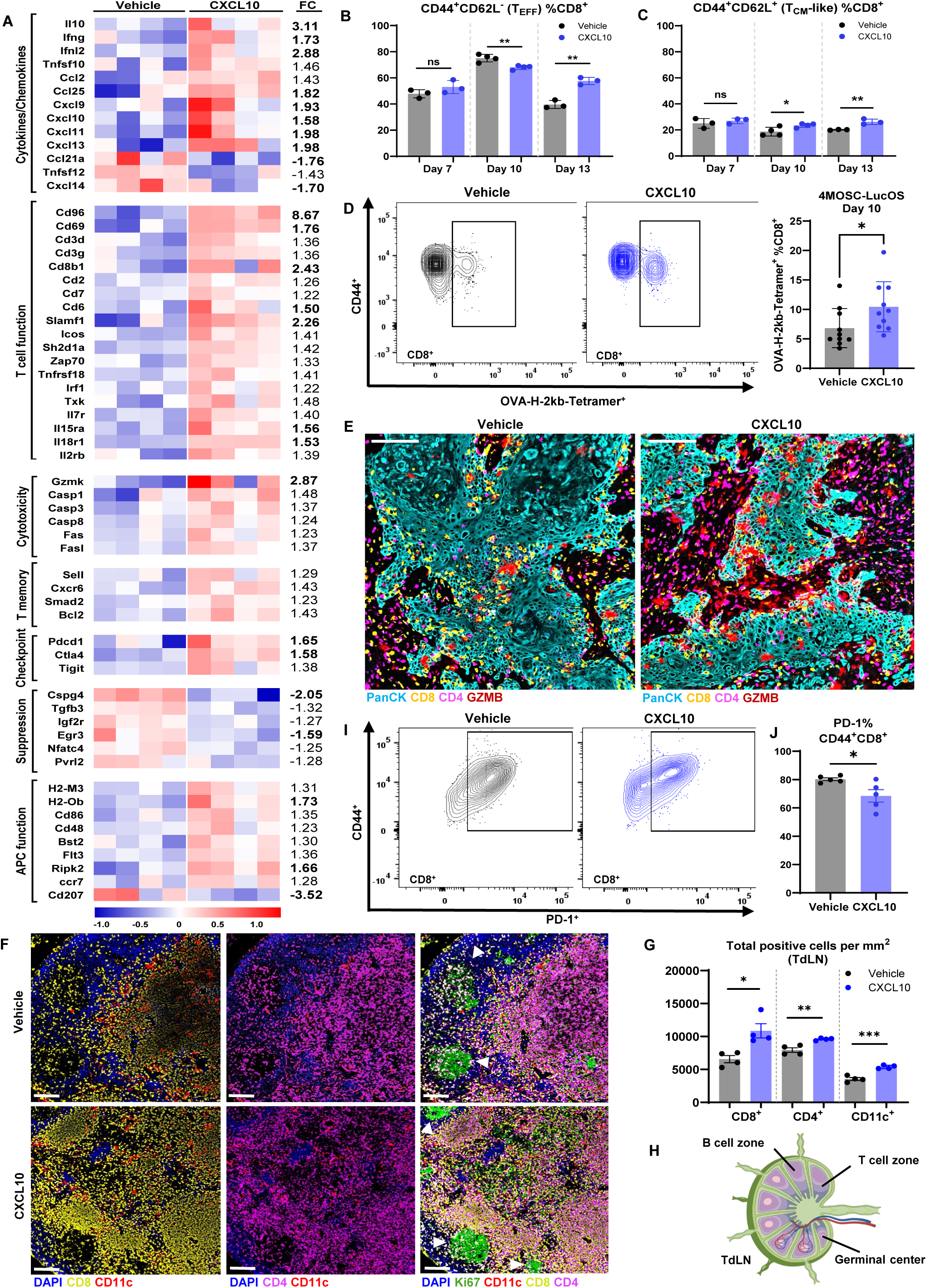
Immune cells isolated from CXCL10-treated tumors exhibit increased activation and tumor antigen-specificity while showing reduced early indications of T cell dysfunction. (**A**) 1.0 x 10^6^ 4MOSC1 cells were implanted into the buccal space of C57BL/6 mice on day 0. Tumors were injected on days 3, 6, and 9 with either vehicle (50 µL of PBS) or 10 µg of CXCL10 in 50 µL of PBS. mRNA from each tumor was isolated and quantified using the NanoString nCounter PanCancer Mouse Immune Profiling gene expression platform. Rosalind software was used to analyze differentially expressed genes (*n* = 4 per group). Differentially expressed genes in vehicle- vs CXCL10-treated 4MOSC1 mice. Fold-change and P value (adjusted) cut-offs are 1.2 (1.5 bolded) and 0.05, respectively. (**B-C**) The frequency of CD44^+^CD62L^-^ cells (T_EFF_) (**B**) or CD44^+^CD62L^+^ cells (T_CM_-Like) (**C**) amongst total CD45^+^CD3^+^ CD8^+^ cells at day 7, 10, and 13 (*n* = 3-4 per group; unpaired Student’s *t* test; data are represented as mean ± SEM). (**D**) Representative flow cytometry plots showing proportion of OVA-H-2kB- Tetramer^+^ cells amongst total CD45^+^CD3^+^ CD8 cells. Right panel, the OVA-H-2kB-Tetramer^+^ cells amongst total CD45^+^CD3^+^CD8^+^ T cells at day 10 (*n* = 10 mice per group; unpaired Student’s *t* test; data are represented as mean ± SEM). (**E**) Representative immunostaining of PanCK (Cyan), CD8 (yellow), CD4 (magenta), and Granzyme B (red) positive cells within the tumors of vehicle and CXCL10-treated mice at day 10. The scale bar represents 100 μm. (**F**) Representative immunostaining of CD8 (yellow), CD4 (magenta), CD11c (red), and Ki67 (green) positive cells within TdLN of vehicle and CXCL10-treated tumors at day 10. The scale bar represents 100 μm. Germinal centers are indicated by ▴. (**G**) Quantification of total CD8+ and CD4+ cells within the TdLN (*n* = 4 mice per group; cell counts from whole TdLN; unpaired Student’s *t* test; data are represented as mean ± SEM). (**H**) Schematic showing the general structure and cell-specific regions within LNs. (**I-J**) Representative flow cytometry plots showing proportion of CD44^+^PD-1^+^ cells amongst total CD45^+^CD3^+^CD8^+^ T cells as well as quantification of these statistics (**J**) at day 10 (*n* = 3-4 mice per group; unpaired Student’s *t* test; data are represented as mean ± SEM).

Results from NanoString gene expression profiling showed that tumors from CXCL10-treated mice not only displayed increased expression of T cell specific genes such as *Cd3d*, *Cd3g*, *Cd7*, and *Cd8b1*, but also many genes associated with T cell activation and cytotoxic function, notably *Cd69*, *Zap70*, *Il18r1*, *Gzmk*, *Fas* and *Fas* ligand (**Fig. 4A**) (*83–87*). One of the most differentially expressed genes amongst treatment groups, *Cd96* (**Fig. 4A**), is known to be highly expressed on effector and memory CD8^+^ T cells as well as on CD4^+^ T effector memory cells (particularly Th1 cells), with expression being particularly elevated on antigen-experienced T cells and tumor-infiltrating lymphocytes (*88, 89*). In addition to an increase in overall infiltration, flow cytometry analysis revealed that a larger proportion of total CD4^+^ and CD8^+^ T cells are CD44^+^CD62L^-^ by day 13 post-engraftment (**Fig. 4B, Fig. S3D**), indicating that CXCL10 treatment favors the formation of T effector (T_EFF_) phenotypes (*90*) that carry out anti-tumoral functions including the secretion of cytotoxic molecules and cytokines. Interestingly, while the total number of CD8^+^ T_EFF_ cells is increased across timepoints, the proportion of CD8^+^ cells displaying a T_EFF_ phenotype is reduced at day 10 within the CXCL10-treated group (**Fig. 4B**). However, this is likely due to an early increase in the proportion of CD44^+^CD62L^+^ cells within the total CD8+ T cells at day 10 that increases through day 13, rather than a larger proportion of naïve-like T cells as seen within vehicle-treated mice (**Fig. 4C**). This suggests the emergence of a prominent population of CD8^+^ T cells with a central memory (T_CM_)-like phenotype (*90, 91*) in the tumors of CXCL10-treated mice. These T_CM_ cells can both self-renew and give rise to a pool of T_EFF_ cells, making them critical for long-term immunity (*91*).

An increase in the expression of many genes involved in the function and maintenance of memory T cell subsets was also revealed from NanoString gene expression profiling (**Fig. 4A**). Notable examples include *Sell* (the gene controlling CD62L expression), *Il7r* (the gene controlling CD127, a marker highly expressed on both T_CM_ and T_EM_ cells) (*92*), *Bcl2* (critical for the persistence of memory T cells) (*93*), and *Cxcr6* (controls the retention of memory T cells) (*94*). Tumors from CXCL10-treated mice also showed a marked increase in *Il10* gene expression (**Fig. 4A**). While there are some indications that IL-10 contributes to an immunosuppressive TME in certain contexts, mice and humans deficient in IL-10 signaling have increased rates of spontaneous tumor formation (*95–97*). IL-10 also enhances the production of IFN-γ and granzyme, facilitates antigen recognition, and is critical for the generation and maintenance memory CD8^+^ T cells, thereby promoting the cytotoxic activity and survival of antigen activated effector and memory CD8^+^ T cells (*98, 99*).

### CXCL10 treatment leads to a greater proportion of tumor antigen-specific CD8^+^ T cells

In addition to promoting effector and memory T cell phenotypes, we hypothesized that CXCL10 treatment would enhance the development of CD8^+^ T cells reactive to tumor-specific antigen (TSA-T cells) that are able to recognize distinct tumor neoantigens and eliminate cancer cells (*100*). To explore this hypothesis, we employed the 4MOSC1-LucOS model in which mice develop 4MOSC1 tumors that express OVA peptide (*67*). Using flow cytometry, we then quantified the abundance of tumor-infiltrating CD8+ TSA-T cells with the OVA-H-2Kb tetramer, as previously described (*67*), and found that tumors from CXCL10-treated mice had a significantly higher proportion of these tumor specific T cells compared to control-treated mice (**Fig. 4D**). To compliment the quantification of TSA-T cells, we evaluated changes in the patterns of GZMB in our 4MOSC1 model in response to CXCL10 since increased release of cytotoxins is indicative of cytotoxic T cell activation and antigen-recognition (*15*). As shown in **Figure 4E**, IF staining revealed more extensive release of GZMB in proximity to CD8^+^ cells infiltrating the tumor parenchyma of CXC10-treated mice.

Presentation of tumor antigens by DCs is a critical step in the activation of cytotoxic CD8+ T cells and their ability to recognize tumor-specific antigens (*25, 66*). Following antigen uptake, DCs undergo maturation, marked by increased expression of costimulatory molecules such as CD80, CD86, and CD40, upregulation of PD-L1, secretion of proinflammatory cytokines, and migration to tumor-draining lymph nodes (TdLN) via CCR7-CCL21α signaling (*101*). Conventional type 1 DCs (cDC1s) also play a pivotal role in regulating the movement of effector T cells into the TME by secreting CXCL9 and CXCL10, indicating crosstalk between cDC1s and T cells within tumors (*102, 103*). Accordingly, we evaluated gene signatures related to DC function and observed a significant increase in those corresponding to costimulatory makers such as *Cd86*, *Cd48*, and *H2-M3,* as well as a key regulator of major histocompatibility complex class I (MHC-I) (*104*), *Nlrc5,* within the tumors of CXCL10-treated mice compared to vehicle controls (**Fig. 4A**) (*105*). A larger proportion of DCs within the tumors of CXCL10-treated mice displayed increased expression of PD-L1 (**Fig. S3G**). Interferons also upregulate MHC-I and II expression and stimulate antigen presentation, with IFN-γ serving as a particularly potent inducer of MHC-II antigen presentation(*106, 107*). Thus, it is notable that tumors from CXCL10-treated mice displayed increased expression of IFN-related gene signatures (*Ifng*, *Ifnl2*, and IFN-inducible CXCR3 chemokines *Cxcl9*, *Cxcl10*, and *Cxcl11*, **Fig. 4A**), as well as many interferon-stimulated genes (*Isi44*, *Ifit1*, *Ifit2*, *Ifit3*, and *Isg15*) (**Fig. S4**). Additionally, there was an increase in *Ccr7* alongside a marked decrease in *Ccl21* expression within the TME following CXCL10 treatment, which may reflect an increase in DC trafficking to the TdLN (**Fig. 4A**). Consistent with the above, tumor rejection following CXCL10 treatment was lost in 4MOSC1 tumor-bearing Batf3^-/-^ mice, which are deficient in cDCs (*108, 109*) (see above, **Fig. 3B**). These data support the dependence of CXCL10-stimulated CD8+ T cells on cDCs for priming to tumor-derived antigens and tumor rejection.

TdLNs are key sites for initiating and amplifying anti-tumoral immune responses, as these critical lymph nodes serve as hubs for antigen presentation by DCs to naïve T cells, inducing rapid proliferation to produce TSA-T cells (*110, 111*). Thus, a higher density of proliferating T cells within the TdLN is indicative of more robust T cell activation, expansion, and trafficking to the TME, and correlates with improved local and systemic immune surveillance for tumor-specific antigen (*111–113*). The specific TdLNs associated with 4MOSC1 buccal tumors have been mapped and are the primary location of T cell priming in the 4MOSC1 model (*67*). Indeed, IF staining of TdLNs from CXCL10-treated mice showed a significant increase in both CD4^+^ and CD8^+^ T cells as well as antigen-presenting cells (CD11c^+^) within the paracortex region (T cell zone) compared to TdLNs from vehicle-treated mice (**Fig. 4F-H**). CD4^+^ and CD8^+^ T cells within the TdLN of CXCL10-treated mice also displayed more robust proliferation, as indicated by increased Ki67 expression (**Fig. 4F**).

Altogether, these data suggest that CXCL10 treatment enhances the recruitment and activation of cells that support cytotoxic function by favoring the formation and maintenance of tumor antigen-specific populations. These results are not only the consequence of CXCL10 impacting T cells, but also indirectly affecting DCs. Moreover, although the treatment is localized to tumors, CXCL10 seems to mobilize a network of interacting immune cells that influence DC and T cell trafficking between tumors and TdLNs.

### CD8+ T cell infiltrating tumors of CXCL10-treated mice exhibit reduced indications of early T cell dysfunction

Without resolution of infection or disease state, effector T cells experience chronic antigen stimulation leading to the progressive loss of effector functions and gradual upregulation of negative costimulatory markers associated with T cell dysfunction (e.g., PD-1^high^) until they reach a terminally exhausted state (TIM-3^high^) (*114*). Thus, one might expect an increase in the expression of PD-1 and other inhibitory molecules (e.g., TIGIT and CTLA4) to be correlated with the increased infiltration of activated TSA-T cells into tumors. This was indeed observed in the tumors of CXCL10-treated mice compared to a vehicle control using the NanoString bulk tumor profiling platform (**Fig. 4A**). However, a significant decrease in PD-1 expression compared to vehicle control was observed when probing only activated CD44^+^CD8^+^ T cell populations by flow cytometry (**Fig. 4I-J**), suggesting an effect of CXCL10 on inhibiting dysfunction. To evaluate if CXCL10 treatment alone can reduce markers of T cell dysfunction outside of the context of the TME, we conducted *in vitro* experiments with primary CD8^+^ T cells from OT-1 transgenic mice whose TCRs are specific for the OVA peptide SIINFEKL (OVA_257–264_) (*115*). Upon stimulation with CXCL10, a decrease in PD-1, CTLA4, and TIGIT expression was observed in CD44^+^CD8^+^ OT-1 T cell populations compared to cells activated with only SIINFEKL peptide (**Fig. S5 A-C**). These results suggest that CXCL10 delays T cell progression towards dysfunction even when antigen stimulation persists.

### IT delivery of CXCL10 angiogenesis and lymphangiogenesis in tumors of 4MOSC1 mice

Beyond its role in leukocyte trafficking, CXCL10 has also been shown to produce anti-angiogenic effects by inhibiting endothelial cell migration and proliferation associated with newly forming lymph and blood vessels in a CXCR3-dependent manner (*116, 117*). Topical application of CXCL10 also blocked both vascular and lymphatic angiogenesis in mouse models of corneal inflammation, preserving vital sensory nerves in the cornea (*118*). We therefore hypothesized that IT delivery of CXCL10 would reduce neovascularization in 4MOSC1 tumors compared to vehicle control. Tumors from CXCL10-treated mice showed downregulation of several genes that promote angiogenesis and play critical roles in regulating epithelial cell adhesion and vascular development, including *DII4* (Delta ligand 4), *Cdh5* (VE-cadherin), and *Vwf* (Von Willebrand factor) (**Fig. 5A**). Elevated expression of Dll4, Cdh5, and Vwf are all recognized as prognostic biomarkers favoring increased metastasis and recurrence in several human cancers (*119–122*). IF staining of 4MOSC1 tumor sections from CXCL10-treated mice also showed decreased lymph and blood vessel density compared to tumors from vehicle-treated mice (**Fig. 5B-C**). This suggests that in addition to recruiting and activating T and NK cells, IT CXCL10 signaling may promote anti-tumoral effects and reduce metastasis by decreasing angiogenesis and lymphangiogenesis within the TME.

**Figure 5.**
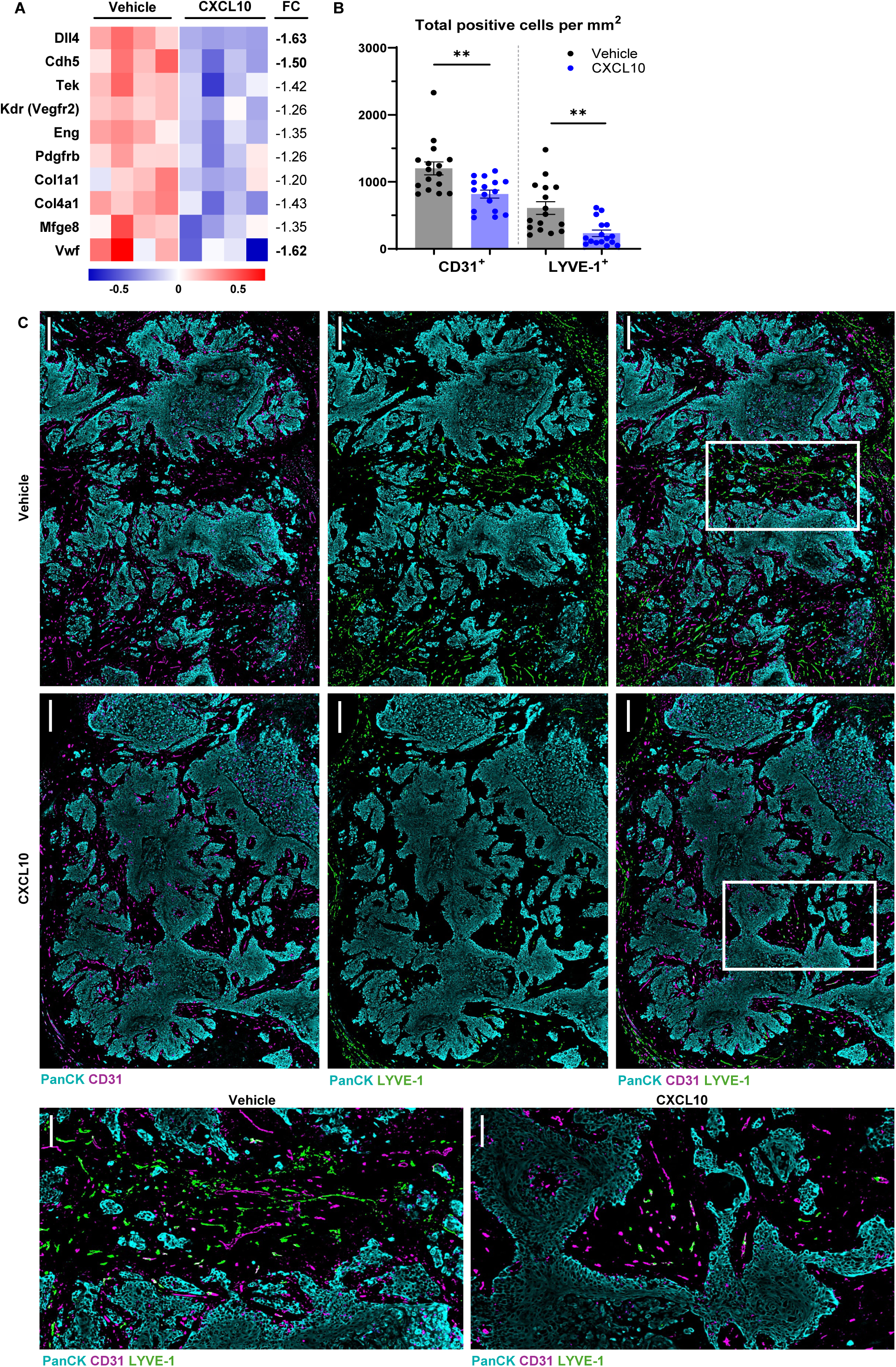
CXCL10-treated tumors exhibit reduced angiogenesis and lymphangiogenesis. (**A**) Differentially expressed genes in vehicle vs CXCL10-treated 4MOSC1 mice. Fold-change and P value (adjusted) cut-offs are 1.2 (1.5 bolded) and 0.05, respectively. (**B**) Quantification of total CD31+ (red) and LYVE-1+ (green) cells within 4MOSC1 tumor sections (*n* = mice per group; 4 ROIs were screened per tumor). (**C**) Representative immunostaining of PanCK (Cyan), CD31 (magenta), and LYVE-1 (green) positive cells of 4MOSC1 tumor sections of vehicle and CXCL10- treated mice at day 10. The scale bar represents 250 μm. The scale bar represents 250 μm. Regions delineated by a white border in the right upper panels are shown at a larger scale in lower panel with a scale bar that represents 100 μm.

### IT CXCL10 synergizes with anti-PD-1 treatment to promote a durable anti-tumoral response in mouse models of HNSCC

When administered as a single agent, CXCL10 inhibited tumor growth and resulted in complete tumor elimination in 24.4% of 4MOSC1 mice on average over all experiments, compared to 18.1% of mice with anti-PD-1 alone (**Fig. S2F**). However, consistent with many molecular therapies, the remaining CXCL10-treated mice developed resistance in subsequent weeks despite initially positive responses, resulting in relapse and tumor progression following the cessation of treatment. Tumor resistance results from the development of immunosuppressive mechanisms that impair anti-tumoral effector functions of CD8^+^ and CD4^+^ T cells, which we showed to be key modulators of the CXCL10 response. It frequently arises from the exploitation of immune checkpoints such as the PD-1/PD-L1 pathway, which plays a central role in regulating T cell activation and exhaustion (*2–4*). PD-1 is predominately expressed on activated T and B lymphocytes, while its ligand PD-L1 is expressed by antigen-presenting cells, such as activated DCs and peripheral blood monocytes, as well as cancer cells to evade immune detection (*4*). Notably, as for human HNSCC, 4MOSC1 tumors characteristically exhibit a high number of infiltrating polymorphonuclear myeloid-derived suppressor cells (PMN-MDSCs) and monocytic myeloid-derived suppressor cells (M-MDSCs) that generally comprise the majority of PD-L1^hi^CD45^+^ immune cells within established tumors (*49, 123*). Thus, the presence of these MDSCs likely contributes to the observed resistance. Indeed, 4MOSC1 tumors from CXCL10-treated mice contained a large percentage of PMN-MDSCs and M-MDSCs that were comparable to the vehicle-treated mice (**Fig. 6A**). However, the PMN-MDSC and M-MDSC populations expressed higher levels of PD-L1 following CXCL10 treatment (**Fig. 6B**), suggesting that the anti-tumor efficacy following CXCL10 treatment might be boosted with the addition of PD-1/PD-L1 blockade to yield more complete responses. To examine the effects of combined therapy in our HNSCC models, anti-PD-1 treatment intraperitoneally (IP) was initiated at the start of the first IT CXCL10 injection (**Fig. 6C**). As shown in **Figure 6D** the combination of CXCL10 and PD-1 blockade resulted in an increase in tumor regression in the 4MOSC1 model, with 40% (**Fig.6D-E, Fig. S2F**; 39.2% on average over all experiments reported in this study) of the mice exhibiting complete and durable responses (>6 months) and, in turn, increased survival when compared to either single agent therapy.

**Figure 6.**
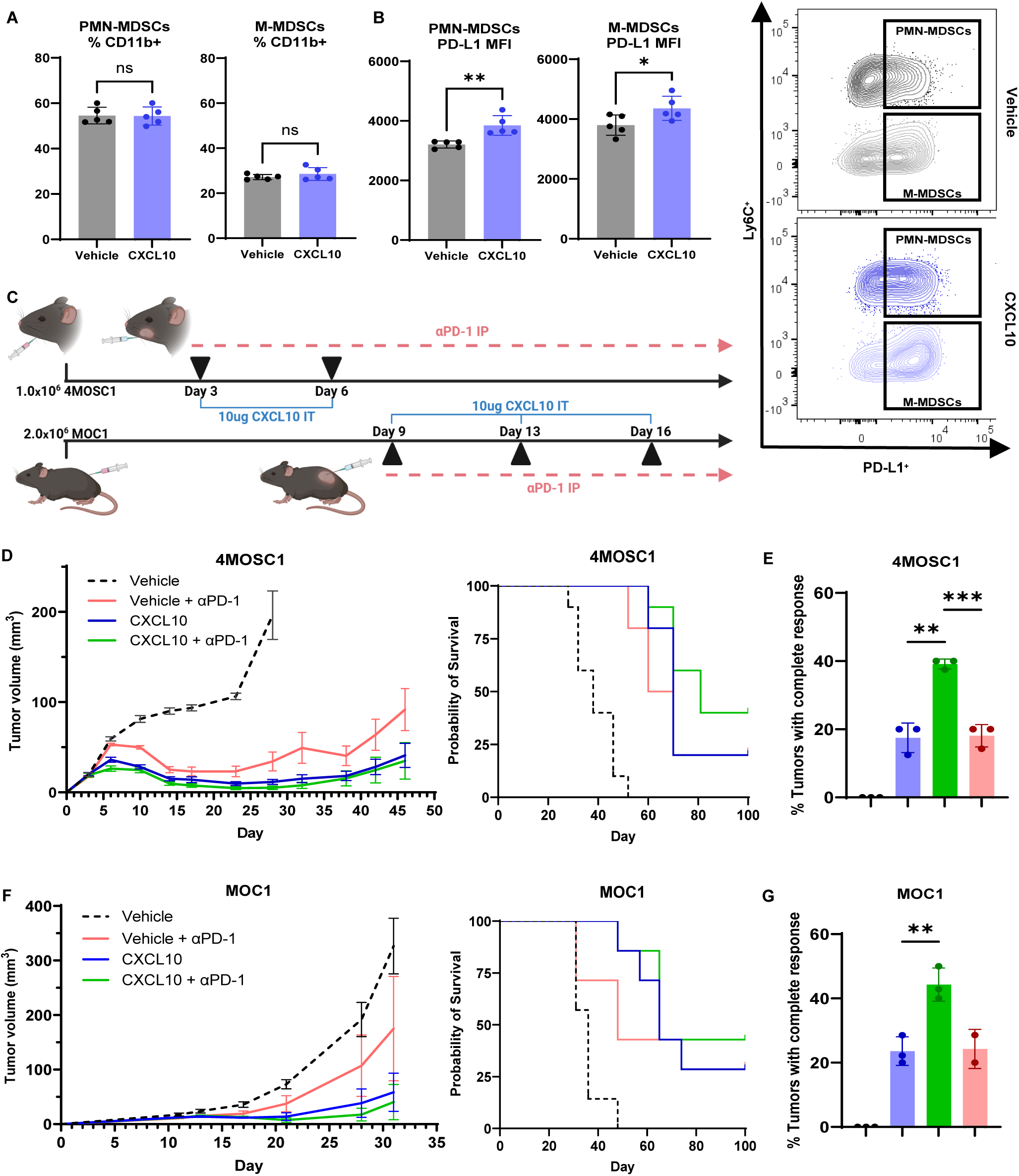
IT CXCL10 synergizes with anti-PD-1 treatment to promote a durable anti-tumoral response. (**A**) The frequency of PMN-MDSC and M-MDSC cells amongst total CD45^+^CD11b^+^ cells (*n* = 5 per group; unpaired Student’s *t* test; data are represented as mean ± SEM). (**B**) The frequency of PD-L1^+^ cells amongst total PMN-MDSC and M-MDSC cells (*n* = 5 per group; unpaired Student’s *t* test; data are represented as mean ± SEM). Representative flow cytometry plots showing relative PD-L1^+^ cells amongst total PMN-MDSC and M-MDSC cells are on the right. (**C**) Schematic detailing experimental setup and dosing schedule for 4MOSC1 and MOC1 models. 1.0 x 10^6^ 4MOSC1 or 2.0 x 10^6^ MOC1 cells were implanted into the buccal space or flank, respectively, of C57BL/6 mice on day 0. Mice received either 10 µg CXCL10 in 50 µL of PBS (per dose) or 50 µL of PBS (vehicle) on the days indicated (▴) and 200 µg of αPD-1 (5 mg/kg) IP every 3 days for 3 weeks initiated at time of vehicle or CXCL10 treatment. (**D, F**) 4MOSC1 (**D**) and MOC1 (**F**) tumor kinetics of mice of indicated treatment groups (*n* = 10 per group (4MOSC1); *n* = 7-8 per group (MOC1)). Survival percentage of 4MOSC1 (**D**) and MOC1 (**F**) mice receiving indicated treatment is shown on the right. (**E, G**) The percent of mice that fully cleared tumors (complete response) for indicated treatment group of matched duplicate experiments. (*n* = 3 per group, with the exception of the αPD-1 treatment group for MOC1 where *n* = 2; unpaired Student’s *t* test; data are represented as mean ± SEM).

Like the 4MOSC1 model, MDSCs are a major immunosuppressive cell population in MOC1 tumors, with PMN-MDSC depletion resulting in the recovery of antigen-specific T cell responses (*124*). Accordingly, the synergistic effect of combining CXCL10 with anti-PD-1 was also explored in the MOC1 model and resulted in 42.9% (**Fig. 6F-G**, **Fig. S2F**; 44.3% on average over all experiments reported in this study) of the mice showing complete tumor clearance and tumor-free survival for 6 months. This is a substantial improvement over treatments with either CXCL10 alone (28.6%, **Fig 6F-G**, **Fig. S2F**; 21.9% on average over all experiments reported in this study) or anti-PD-1 alone (28.6%, **Fig 6F-G**, **Fig. S2F**; 20.54% on average over all experiments reported in this study).

## DISCUSSION

The successful infiltration of cytotoxic effector cells remains a critical determinant of the efficacy of ICBs such as PD-1/PD-L1 blocking agents. The CXCR3-CXCL10 chemokine system has demonstrated functional relevance in both the successful recruitment of CD8+ T cells into the TME, as well as enhancing the proliferation and function of intratumoral CD8+ T cells during anti-PD-1 treatment (*27*). In fact, studies using the CERTIM pan-cancer cohort indicate that high expression of CXCR3 and its ligands were amongst the top correlative indications of positive responses to ICB therapy (*125*). In this study, we demonstrated that intratumoral augmentation of CXCL10 increases the infiltration of cytotoxic effector cells, leading to suppressed tumor growth and improved outcomes in murine models of HNSCC. This included enhanced infiltration of CD8+ T cells, CD4+ T cells, and NK cells, all of which contributed to tumor regression in CXCL10-treated mice.

However, our data suggests that IT CXCL10 drives tumor clearance not only though boosting intratumoral infiltration of cytotoxic immune cells and cells that support cytotoxic function, but also by favoring conditions that promote the effector function of T cells. Previous studies have shown that CXCL10 induces CD4+ T cell polarization towards immune-supporting Th1 and Th17 lineages via STAT1, STAT4, and STAT5-dependent signaling (*80, 81*). Conversely, CXCL11 causes CD4+ T cell polarization towards the immunosuppressive type 1 regulatory (Tr1) lineage (IL-10^high^) mediated by STAT3- and STAT6-dependent pathways (*38*). Additional data indicate that CXCL10 promotes the terminal differentiation of effector CD8+ T cells within the spleen (*82*) and facilitates T cell interactions with DCs (*39*). CXCL10 has also been shown to stimulate PD-L1 expression on NK cells, enhancing the ability of these cells to stimulate the proliferation of T cells and increase production of IFN-γ and TNFα *in vitro* (*126*). Together, these data suggest that CXCL10 not only acts as a chemoattractant but also influences the biological responses of CXCR3-expressing immune cells, potentially having a direct influence over their maturation and differentiation. Here, we showed that T cells recruited to the TME of CXCL10-treated mice displayed increased tumor antigen specificity, penetrance into the tumor parenchyma, and evidence of cytotoxic effector function, such as granzyme release.

Following priming with tumor-derived antigen, a subset of naïve T cells (T_N_) proliferate, upregulate markers of activation, and downregulate circulatory markers like CD62L, differentiating into CD44^+^CD62L^-^ effector T cells (T_EFF_) that carry out anti-tumoral functions including the secretion of cytotoxic molecules and cytokines. Other primed T_N_ cells differentiate into memory T cells, including long-lived central memory T cells (T_CM_), characterized as CD44^+^CD62L^+^, as well as stem (T_SCM_), effector (T_EM_), and tissue residential (T_RM_) memory T cell subsets (*64, 90*). T_CM_’s in particular can both self-renew and give rise to a pool of T_EFF_, making them critical for long-term immunity. Our studies revealed an increase of T cells displaying T_CM_-like markers in response to CXCL10 treatment, also suggesting that CXCR3-signaling may favor the formation of memory T cell subsets.

Our data also suggests that CXCL10 signaling through CXCR3 directly slows progression to dysfunction. Persistent antigen exposure can drive T cell dysfunction and exhaustion, characterized by reduced effector function and sustained expression of inhibitory receptors (*23, 114*). In contrast, timely antigen clearance promotes the development of memory T cells. Thus, the duration and intensity of antigen signaling determines, in part, whether T cells become part of the functional memory pool or succumb to the progressive dysfunction associated with exhaustion. However, T cell dysfunction is not solely regulated by the persistence of antigen. Recent work demonstrated that prostaglandin E2 (*PGE2*) contributes to T cell dysfunction through the stimulation of Gα_s_/cAMP signaling pathways in models of chronic stimulation of primary T cells *in vitro* (*115*). Using the same model, CXCL10 treatment was shown to rescue T cells from dysfunction as reflected by restored levels of IFN-y and TNFα in the presence of PGE2 (*115*). The mechanism is likely through CXCR3-mediated stimulation of Gα_i_, which counters Gα_s_/cAMP activation by PGE2. Consistent with these data, it has been demonstrated that CXCR3^−^ PD-1^hi^ CD8^+^ T cells express higher levels of inhibitory molecules (including TIM-3, LAG-3 and TIGIT) than the other subsets, indicating a higher level of T cell dysfunction (*27*).

The mechanisms by which CXCL10 treatment inhibits tumor growth do not appear limited to CXCR3-expressing immune cells. Consistent with the increase in TSA-T cells that accumulate in 4MOSC1 tumors in response to CXCL10 treatment, cDCs were shown to play a critical role in the efficacy of the CXCL10 treatment. We also observed increased antigen-presenting (CD11c^+^) cells in TdLNs where these cells prime and activate CD8+ and CD4+ T cells, which also accumulate in the TdLNs. This is striking because administration of CXCL10 into tumors appears to promote the trafficking of CXCR3- and non-CXCR3-expressing cells between tumors and TdLNs, and the establishment of a network of anti-tumoral cells that results in the elimination of tumor cells.

Finally, CXCL10 does not just affect immune cells. We also observed a dramatic reduction of both angiogenesis and lymphangiogenesis in tumors from CXCL10-treated mice compared to control mice, consistent with prior reports and implicating CXCR3 signaling on endothelial cells (*116, 118*). These data further define the multimodal mechanisms for the efficacy of CXCL10 treatment, as it is well established that increased rates of angiogenesis in solid tumors is essential for the maintenance of tumor growth, progression, and distal metastasis (*21, 117*). Additionally, excessive angiogenesis produces a vasculature that is both structurally and functionally abnormal, reducing vasculature normalization, limiting blood perfusion, and interfering with the infiltration of immune cells (*21*). Inhibition of angiogenesis also enhances T-cell infiltration and drug delivery to tumors, which further enhances the efficacy of ICB treatments (*127*).

Importantly, IT delivery of CXCL10 as a single agent was sufficient to achieve significant anti-tumoral responses in two syngeneic murine models of HNSCC, with the rate of complete tumor clearance rivaling that of anti-PD-1 therapy alone at ∼24.4%. However, it has been reported that responsiveness to anti-PD-1 therapy is *dependent* on CXCR3 and its chemokines, as CXCR3, CXCL9 and CXCL10 knockout mice exhibit markedly reduced survival in the anti-PD-1 responsive MC38 tumor model (*27*). Indeed, complete tumor remittance increased from 18-24% with CXCL10 and anti-PD-1 as single agents to 39.2% and 44.3% in 4MOSC1 and MOC1 models, respectively, when given in combination (**Fig. S2F**). This synergistic anti-tumoral efficacy is consistent with prior studies of anti-PD-1 responsive MC38 colon adenocarcinoma and MCA1956 fibrosarcoma tumor models, which exhibited significantly higher levels of CXCL9 and CXCL10 than anti-PD-1 resistant tumors (*27*).

As immune modulators that are generally administered systemically, ICBs such as anti-PD-1 and anti-CTLA-4 also disrupt normal immune regulation and tolerance in healthy tissues (*128, 129*). The resulting unbridled immune activity can induce a wide spectrum of immune-related adverse events (irAEs) such as dermatitis, colitis, thyroiditis, and pneumonitis in a significant proportion of patients (*128–130*). CTLA-4 blocking agents like ipilimumab have particularly high rates of irAEs, which are reported in up to 60-70% of patients compared to 30% in those treated with anti-PD-1 (*128*). Thus, in addition to improving the efficacy of ICBs, it will be important to determine if CXCL10/ICB combination treatments reduce the risk of irAEs by enabling anti-tumor efficacy to be achieved with a lower dose of the ICB. Meanwhile, localized therapeutics like CXCL10 may inherently have a reduced risk of irAEs by limiting chemokine activity to the tumor and minimizing systemic exposure and off-target immune activation in healthy tissues. As increased localized CXCL10 levels produced a higher proportion of TSA-T cells in our studies, off-target immune cell overactivation is expected to be minimized due to a more tumor-restricted immune response.

Our results are consistent with prior studies using epigenetic modulators and vaccination via CXCL9/CXCL10-engineered DCs; these studies demonstrated that amplification of CXCR3 signaling within the TME leads to improved responses to ICB in models of colorectal carcinoma, melanoma, and non-small cell lung cancer (*27, 131*). mRNA-based therapeutics are also emerging as a platform for the intratumoral delivery of other immunomodulatory proteins (*132*). While promising, these methods are less controlled with respect to CXCL10 levels and duration of treatment, which could impact efficacy and undesired effects. Our approach with intratumoral delivery of recombinant chemokine offers a more immediate and controlled therapeutic approach that can be discontinued or adjusted as needed while presenting a lower risk of immunogenicity and off-target activity. However, while significant efficacy was observed in the HNSCC models, responses to CXCL10 as a single agent as well as in combination with anti-PD-1 were still limited, as tumors regrew in ∼75% and ∼54-60% of the mice, respectively. Thus, additional modulators such as inhibitors of MDSCs, which are abundant in the HNSCC TME, may be worth exploring to determine if it is possible to further sensitize tumors to CXCL10-therapy. Consistent with this, recent work has demonstrated that hydrogel delivery of PD-L1 and inhibitors of DPP4 (a protease that rapidly deactivates CXCR3 ligands) greatly reduced degradation of CXCL10, promoting cancer regression in a lung metastasis model (*133*). Finally, the use of CXCL10 as an immune-stimulating molecule may also be applicable to other cancer models beyond HNSCC, particularly those also characterized by an immune suppressive tumor microenvironment.

Taken together, our study shows that IT CXCL10 is a viable approach for enhancing the intratumoral recruitment of CXCR3 cells into the TME, as well as for promoting their effector functions, differentiation into long-term memory cells, and resistance to dysfunction. CXCL10 also appears to stimulate a larger network of interacting cells that are critical for tumor elimination and reshaping the vasculature to a state that is unfavorable for tumor proliferation and metastasis. Our findings provide the rationale for the future clinical evaluation of IT CXCL10 as a multimodal strategy to overcome resistance to ICB immunotherapies and improve outcomes in HNSCC patients and possibly other cancers.

## MATERIALS AND METHODS

### CXCL10 production and purification

A previously described *E. coli* expression/purification system involving chemokine fused to the C-terminus of a His-tagged ubiquitin variant was used to produce murine CXCL10; the variant contains a tryptophan-serine sequence at the C-terminus to facilitate accurate determination of concentration (*68*). The His-tag enables Ni-affinity chromatography purification of the detergent solubilized fusion protein, which is subsequently refolded in a redox buffer to form the correct disulfides. The ubiquitin facilitates cleavage of CXCL10 from ubiquitin by the deubiquitylating enzyme Usp2cc, leaving the native chemokine N-terminus that is required for WT activity. A final purification step by reversed phase HPLC is used to isolate the desired protein with purity suitable for *in vivo* use and a yield of ∼15 mg/L. The identity of the final product was confirmed by mass spectrometry and the purity assessed via SDS-PAGE gel. The protein sequence of the final product:

### IPLARTVRCNCIHIDDGPVRMRAIGKLEIIPASLSCPRVEIIATMKKNDEQRCLNPESKTIKNLMKAFSKRSKRAPWS

Functional integrity was assessed by cell migration assays with CXCR3^+^ Jurkat cells (kind gift from Sudar Rajagopal, Duke University) and bioluminescent resonance energy transfer (BRET)-based G protein activation and β-arrestin recruitment assays (*134, 135*).

### Cell lines

The 4MOSC1 cell line was generated in-house (*49*). MOC1 cells were generously provided by Dr. R. Uppaluri (Harvard Medical School, Boston, MA) (*50*). Authentication of cell lines was confirmed by multiplex STR profiling (Genetica DNA Laboratories, Inc. Burlington, NC) to ensure cell identity. 4MOSC1 cells were grown on culture dishes pre-coated with collagen (Corning, #354249) in Defined Keratinocyte-SFM medium (#10744019; ThermoFisher) supplemented with 5 ng/mL EGF Recombinant Mouse Protein (#PMG8044; ThermoFisher), Defined Keratinocyte SFM Growth Supplement (1 vial/500 mL media) (#10784015; ThermoFisher), 50 pM Cholera Toxin (#C8052; Sigma-Aldrich), and 1% antibiotic/antimycotic solution (#BW17-603E; Fisher). MOC1 cells were cultured in a 2:1 IMDM (#12440-046; ThermoFisher) and Hams Nutrient Mixture (#11765054; ThermoFisher) supplemented with 5% fetal bovine serum (# sh30071.03; HYCLONE), 1% antibiotic/antimycotic solution (#BW17-603E; ThermoFisher), 5 mg/mL insulin (#10516; Sigma-Aldrich), 400 ng/mL Hydrocortisone (# H0135; Sigma-Aldrich,), and 5 ng/mL EGF (#01-107; Millipore). All cells were cultured at 37°C in the presence of 5% CO_2_.

### Antibodies

PD-1 antibody (clone J43, BE0033-2), isotype antibody (Armenian hamster IgG isotype control, BE0091), and CD8 depletion antibody (clone YTS 169.4, BE0117), CD4 depletion antibody (clone GK1.5, BE0003-1), and NK1.1 depletion antibody (clone PK136, BE0036) were obtained from Bio X Cell. Fluorochrome-conjugated antibodies were purchased from BD Biosciences (San Jose, CA) and BioLegend (San Diego, CA) unless otherwise indicated.

### Mice

All the animal studies using HNSCC tumor xenografts and orthotropic implantation studies were approved by the University of California, San Diego Institutional Animal Care and Use Committee (IACUC), and performed according to protocol ASP #S15195, with all experiments adhering to relevant ethical regulations for animal testing and research. Mice were housed at the Moores Cancer Center (University of California, San Diego) in a Micro-Isolator and individually ventilated cages supplied with acidified water. The mice were fed 5053 irradiated Picolab rodent diet 20 from LabDiet. The temperature in the facility was maintained between ∼18 and 23°C, with 40– 60% humidity, and a 12 h light/12 h dark cycle. All animal manipulation activities were conducted in laminar flow hoods within the facility. Wildtype C57BL/6N mice were obtained from Charles River Labs. CXCR3^-/-^ mice (B6.129P2-Cxcr3tm1Dgen/J), BATF3^-/-^ mice (Batf3tm1Kmm/J), and OT-1 mice (C57BL/6-Tg(*TcraTcrb*)1100Mjb/J, 003831) were obtained from the Jackson Laboratory (Bar Harbor, ME) and bred in-house.

### In vivo mouse tumor models and analysis

The ARRIVE essential 10 guidelines (https://arriveguidelines.org/arrive-guidelines) were followed to conduct all *in vivo* animal studies (murine). For the MOC1 model, 2 million MOC1 cells were implanted into the right flanks of female C57BL/6N or CXCR3^-/-^ mice (4–6 weeks of age). When average tumor volumes reached 40–60 mm^3^, (around day 10), the mice were randomized into groups for subsequent experiments. For the 4MOSC1 model, 1 million cells were transplanted into the right buccal space of female C57BL/6N, CXCR3^-/-^, or BATF3^-/-^ mice (4–6 weeks of age). When average tumor volumes reached 10-20 mm^3^ (around day 3), the mice were randomized into groups and treatment initiated. For IT treatment, 10 µg chemokine in 50 µl PBS or vehicle control (50 µl PBS) were injected directly into tumors. 4MOSC1 mice were treated on days 3 and 6 post tumor cell implantation, and occasionally received a 3^rd^ dose on day 9, as indicated in the individual experiments. MOC1 mice were treated three times (days 10, 14, and 17). For studies with checkpoint blockade inhibitors, the mice were treated IP with 10 mg/kg anti-PD-1 antibody (clone J43) three times a week for 3 weeks. For studies with depletion antibodies, mice were treated IP with 10 mg/kg of the indicated depletion antibody three times a week for 3 weeks. Tumor volumes were calculated as (width^2^)(length)/2 (mm^3^). The mice were euthanized after completion of the treatment or when control-treated mice succumbed to tumor burden. As needed, tumors were dissected for flow cytometric analysis or histologic and immunohistochemical evaluation. The GraphPad Prism version 10 statistical analysis program (GraphPad Software, Boston, MA) was used to produce figures.

### Flow cytometry

Flow cytometry using comprehensive immune cell panels were utilized to gauge changes in immune cell infiltration and activation status in vehicle vs CXCL10-treated 4MOSC1 mice. Tumors were isolated, minced, and resuspended using the Tumor Dissociation Kit (#xxx; Miltenyi Biotec, San Diego, CA), diluted into DMEM for subsequent processing with the gentleMACS Octo Dissociator. Digested tissues were then passed through 70-µm strainers to produce a single-cell suspension. Samples were washed with PBS and processed for live/dead cell discrimination using Zombie viability stain (BioLegend, San Diego, CA). Cell suspensions were then washed with cell staining buffer (#420201; BioLegend) prior to staining with antibodies for 30 min at 4°C, protected from light. The following mouse antibodies and dilutions were used: CD45 (30-F11, 1:100), CD3 (17A2, 1:200), CD8a (53-6.7, 1:100), CD4 (RM4-4, 1:100), Slamf6 (330AJ, 1:100), PD-1 (29 F.1A12,1:100), CD44 (IM7, 1:100), CD19 (6D5, 1:100), CXCR3 (S18001A, 1:100), Tim3 (RMT3-23, 1:100), NK1.1 (PK136, 1:100), CD69 (H1.2F3, 1:100), CD62L (MEL-14, 1:100), Ly6C (HK1.4, 1:100), CD11b (M1/70, 1:100), CD11c (N418, 1:100), XCR1 (ZET, 1:100), CD64 (X54-5/7.1, 1:100), CD103 (2E7, 1:100), MHCII (M5/114.15.2, 1:200), CD80 (16-10A1, 1:100), IFNγ (XMGl.2, 1:100), granzyme B (GBll, 1:100), TNF-α (MP6-XT22, 1:100), LAG3 (C9B7W, 1:100), and Ki-67 (16A8, 1:100). Stained cells were washed and then fixed with BD cytofix for 20 min at 4°C, protected from light. In the case of intracellular staining, permeabilization was performed following surface staining by incubating the cells with intracellular targeted antibodies in fixation-permeabilization buffer (#88-8824-00; ThermoFisher) for 30 min at 4°C and protected from light. Samples were analyzed using a BD LSR Fortessa or BD LSR Fortessa X-20. Downstream analysis was performed using FlowJo version 10.8.1 (BD Life Sciences). The gating strategy for identifying NK, CD4, and CD8 T cell subsets is shown in **figure S3C**. The GraphPad Prism version 10 statistical analysis program (GraphPad Software, Boston, MA) was used to produce figures.

### T cell activation with CXCL10 *in vitro*

For T cell activation studies, a protocol was adapted from (*115*). Briefly, splenocytes were isolated from 4- to 6-week-old mice and mechanically disrupted. Red blood cells were lysed in red blood cell lysis buffer (BioLegend) according to manufacturer’s instructions. CD8^+^ T cells were isolated with an EasySep CD8 isolation kit by negative selection. For initial activation, naïve CD8+ T cells were cultured (0.5-1 million cells per mL) in 24-well plates with 50 U mL^−1^ IL-2, 1 µg/mL of anti-mouse CD3 (#100340; BioLegend), and 2 µg/mL of anti-mouse CD28 (#102116; BioLegend) for 48 h. Cells were then collected, replated, and cultured in either 50 U/mL IL-2 (PeproTech) alone, 50 U/mL IL-2 with 100 nM OVA_257–264_ peptide, or 50 U/mL IL-2 with OVA_257–264_ and CXCL10 (concentrations varying from 1-100 nM) for 24-48 h. Cells were then washed with cell staining buffer and both surface and intracellular staining was performed as previously described above with the following antibodies: CD45 (30-F11, 1:100), CD3 (17A2, 1:200), CD8a (53-6.7, 1:100), CD4 (RM4-4, 1:100), Slamf6 (330AJ, 1:100), PD-1 (29 F.1A12, 1:100), CD44 (IM7, 1:100), CXCR3 (S18001A, 1:100), Tim3 (RMT3-23, 1:100), CD62L (MEL-14, 1:100), IFNγ (XMGl.2, 1:100), granzyme B (GBll, 1:100), TNF-α (MP6-XT22, 1:100), LAG3 (C9B7W, 1:100), and Ki-67 (16A8, 1:100). Stained cells were washed and then fixed with BD cytofix. For intracellular and transcription factor staining, cells were stimulated with 1X cell activation cocktail with brefeldin A (BioLegend) in medium for 4–6 h at 37 °C before viability staining; unstimulated cells served as control. After surface staining, cells were fixed with a FOXP3/transcription factor buffer set (Invitrogen) and stained with intracellular antibodies for 45 min at room temperature. Samples were then analyzed using a BD LSR Fortessa or BD LSR Fortessa X-20. Downstream analysis was performed using FlowJo version 10.8.1 (BD Life Sciences). The GraphPad Prism version 10 statistical analysis program (GraphPad Software, Boston, MA) was used to produce figures.

### Quantification of antigen-specific T cells *in vivo*

To detect TSA-T cells in the TME, 1 million 4MOSC1-LucOS cells, generation described in Saddawi-Konefka et al., 2022, were implanted into the right buccal space of female C57Bl/6 mice. When the average tumor volume reached 10-20 mm^3^, mice were randomized into groups and treatment was initiated. Treatments included 10 µg chemokine in 50 µl of PBS or 50 µl PBS (vehicle control) which were injected directly into tumors on days 3 and 6. At days 10-15, tumors were isolated and processed to obtain a single-cell suspension as described above. For antigen-specific T cell tetramer staining, Flex-T™ Biotin H-2 K(b) OVA Monomer (#280051; BioLegend) paired with PE-streptavidin or APC-streptavidin (#405203 or 405207, respectively; BioLegend) were used according to manufacturers’ instructions and as previously described (*67*). Samples were analyzed using a BD LSR Fortessa or BD LSR Fortessa X-20. Data analysis was performed using FlowJo version 10.8.1 (BD Life Sciences), version 10.8.1. The GraphPad Prism version 10 statistical analysis program (GraphPad Software, Boston, MA) was used to produce figures.

### The Cancer Genome Atlas (TCGA) analysis

The cBio Portal (http://www.cbioportal.org/) was utilized to download and analyze transcriptomic data from the TCGA PanCancer Atlas data set for HNSCC (*51, 52*). Hazard ratios were calculated from the log-rank test between cohorts to determine the relationship between CXCR3 and CXCL10 on overall outcomes and inflammatory status in HNSCC patients. CXCR3^high^/CXCL10^high^ cohorts (n=128 and 132) and CXCL10^low^/CXCL10^low^ cohort (n=131 and 126) represent the upper 25% and lower 25% threshold for *Cxcr3* or *Cxcl10* mRNA expression respectively. Differentially expressed genes displayed in figures represent Log_2_ fold-change and P value (adjusted) cut-offs are 1 and 0.05, respectively between CXCL10^high^ vs CXCL10^low^ cohorts using Wilcoxon Rank Sum test. Average mRNA gene expression amongst CXCL10^high^ vs CXCL10^low^ cohorts were used to generate cell enrichment scores using the xCell Cell Enrichment (https://xcell.ucsf.edu/) analysis platform (*63*). The GraphPad Prism version 10 statistical analysis program (GraphPad Software, Boston, MA) was used to produce figures.

### NanoString nCounter Platform

RNA was isolated by homogenization of whole tumors in TRIzol® (Invitrogen) followed by phenol:chloroform extraction and RNeasy Mini Kt based column purification with on-column DNase treatment (Qiagen). Hybridization of samples was performed at The Next Generation Sequencing Core (Salk Institute for Biological Studies, La Jolla, CA) according to the NanoString Hybridization Protocol for nCounter XT CodeSet Gene Expression Assays. Samples were run on the nCounter SPRINT Profiler with the nCounter PanCancer Mouse Immune Profiling gene expression platform. Data was analyzed by ROSALIND® (https://rosalind.bio/) (San Diego, CA). Read Distribution percentages, violin plots, identity heatmaps, and sample MDS plots were generated as part of the QC step. Normalization, fold changes and p-values were calculated using criteria provided by NanoString. ROSALIND® follows the nCounter® Advanced Analysis protocol of dividing counts within a lane by the geometric mean of the normalization probes from the same lane. Housekeeping probes used for normalization were selected based on the geNorm algorithm as implemented in the NormqPCR R library (Perkins et al., 2012). Abundance of various cell populations was calculated using the Nanostring Cell Type Profiling Module with ROSALIND software. ROSALIND performs a filtering of Cell Type Profiling results to include results that have scores with a p-Value greater than or equal to 0.05. Fold changes and pValues were calculated using the fast method as described in the [nCounter® Advanced Analysis 2.0 User Manual](https://www.nanostring.com/download_file/view/1169/3842). P-value adjustment was performed using the Benjamini-Hochberg method of estimating false discovery rates (FDR). Clustering of genes for the final heatmap of differentially expressed genes was done using the PAM (Partitioning Around Medoids) method using the fpc R library (*136*) that takes into consideration the direction and type of all signals on a pathway, the position, role and type of every gene, etc. Hypergeometric distribution was used to analyze the enrichment of pathways, gene ontology, domain structure, and other ontologies. The top GO R library was used to determine local similarities and dependencies between GO terms in order to perform Elim pruning correction. Several database sources were referenced for enrichment analysis, including Interpro (*137*), NCBI (*138*), MSigDB (*139, 140*), REACTOME (*141*), WikiPathways (*142*). Enrichment was calculated relative to a set of background genes relevant for the experiment. GraphPad Prism version 10 statistical analysis program (GraphPad Software, Boston, MA) was used to produce figures.

### Microscope image acquisition

Tissues (buccal tumors and cervical lymph nodes) were harvested, fixed in zinc formalin fixative (Sigma-Aldrich) and sent to the Biorepository and Tissue Technology Shared Resources (BTTSR) at Moores Cancer Center (San Diego, CA) for paraffin embedding, sectioning, and IF staining. Histology samples were analyzed using QuPath 0.5.1 (Edinburgh, UK). The following antibodies were used according to the BTTSR core’s protocol: CD4 (anti-mouse, Abcam), CD8 (anti-mouse, Invitrogen), CD19 (anti-mouse, Cell Signaling), CD25 (anti-mouse, Cell Signaling), PanCK (anti-mouse, Dako), Granzyme B (anti-mouse, Invitrogen), CD11c (anti-mouse, Cell Signaling), CD80 (anti-mouse, Abcam), NK1.1 (anti-mouse, Cell Signaling), Ki67 (anti-mouse, GeneTex), LYVE-1 (anti-mouse, Invitrogen), and PD-1 (anti-mouse, Abcam). Slides were scanned using a PhenoImager HT (Akoya Biosciences) at 40x resolution. H&E stained sections underwent brightfield whole slide scanning using an AT2 Aperio (Leica Biosystems) at 40x resolution. All image analyses were performed using the QuPath software to perform pixel classification of stained cells and positive cell counts.

### Statistical analysis

The sample size for each experiment was selected based on historical data and previous publications using the same tumor models (PMID: 31804466, PMID: 35879302). Mice from *in vivo* experiments were randomized before initiation of treatment or data collection. Data collection and analysis were not performed blind to the conditions of the experiments. Data analyses, variation estimation, and validation of test assumptions were conducted with GraphPad Prism version 10 statistical analysis program (GraphPad Software, Boston, MA) unless otherwise indicated. Differences between the tumor kinetics amongst experimental groups were analyzed using simple linear regression analysis. The differences between experimental groups in cell density, response values, and tumor volume were analyzed using independent *t*-tests or ANOVA (when >2 conditions). Survival analysis was performed using the Kaplan–Meier method and log-rank tests. The asterisks in each figure denote statistical significance, or ns for non-significant *p* > 0.05; \**p* < 0.05; \*\**p* < 0.01; and \*\*\**p* < 0.001. All the data are reported as mean ± SEM (standard error of the mean). Experiments were independently repeated at least three times with similar results.

### Supplemental material summary

Fig. S1 lists the top 100 differentially expressed genes between CXCL10^high^ and CXCL10^low^ HNSCC tumors. **Fig. S2 A-E** shows the production and functional verification of produced CXCR3 ligands. **Fig. S2 F** shows a table detailing the proportion of mice exhibiting complete tumor clearance across multiple independent experiments for tumors treated with indicated treatment. **Fig. S3 A-B** shows representative IF images and quantification of NK cells within 4MOSC1 tumors of vehicle vs CXCL10 treated mice. **Fig. S3 C-H** shows supplemental flow data as referenced in the main text along with a representative gating strategy for T cells. **Fig. S4** shows an extended list of differentially expressed genes from 4MSOC1 tumors of vehicle and CXCL10 treated mice. **Fig. S5** shows that CXCL10 treatment promotes proliferation and reduced inhibitory markers in primary OT-1 CD8+ T cells *in vitro*.

## DATA AVAILABILITY

(NOTE: Data files are currently being uploaded into detailed archives at the time of this submission and can be available from the corresponding author upon request for review)

The final analyzed tumor growth kinetic data and protocols will be made available at the time of associated publication, with all raw data (.xlsx), analyzed data (.prism) and metadata will be uploaded to the public repository Mendeley Data (https://data.mendeley.com/) [persistent link to data in repository, e.g., DOI, accession numberXXXX]. For images, all raw data (.xlsx), analyzed data (.prism) and metadata will be uploaded to the public repository BioImage Archive (https://www.ebi.ac.uk/bioimage-archive/). Flow flow cytometry data, all raw data (.fcs), analyzed data (.flowjo) and metadata will be collected and annotated according to the MIFlowCyt standard and uploaded to the FLOWrepository (www.flowrepository.org) [persistent link to data in repository, e.g., DOI, accession numberXXXX]. The final analyzed NanoString data and protocols will be made available at the time of associated publication, with all raw data (.rcc), analyzed data (.xlsx) and metadata will be uploaded to the NCBI-supported public functional genomics data repository Gene Expression Omnibus (GEO) (https://www.ncbi.nlm.nih.gov/geo/). [persistent link to data in repository, e.g., DOI, accession numberXXXX]. All additional data that support the findings in this study are available from the corresponding author upon request.

## ACKNOWLEDGEMENTS

This project was supported by grants from the National Institute of Dental and Craniofacial Research (NIH/NIDCR R01DE033909-01) and the Moores Cancer Center, Delivering Discoveries: Jawsome Shark Tank Multidisciplinary Pilot Project Program (NIH 2P30CA023100), both to T.M. Handel. and J.S. Gutkind. C. Shinn was supported by the Pharmacological Sciences Pre-Doctoral Training Program (5T32GM007752-40), the Tribal Member Initiative (TMI) Pre-doctoral Fellowship, and the Cancer Cell Signaling and Communication Post-doctoral Training Grant (NIH/NCI T32 CA009523). R. Saddawi-Konefka (UCSD) was supported by an NRSA Training Award (NIH/NIDCR F32DE029990-01). We would like to acknowledge Elsa Molina at the Salk OMICS core for the preparation and running of samples with the nCounter PanCancer Mouse Immune Profiling gene expression platform (NanoString Technologies) and training on data as well as Cheryl Kim at the LJI Flow Cytometry Core and Michael Rose and Valeria Estrada from the UCSD Biorepository and Tissue Technology Shared Resource Core. Finally, we would like to thank Dr. John Chang and Dr. Jack Bui (UCSD) for helpful discussions. All cartoon renderings were created using https://www.biorender.com (BioRender.com).

## REFERENCES

1. J. V. Poulose, C. T. Kainickal, Immune checkpoint inhibitors in head and neck squamous cell carcinoma: A systematic review of phase-3 clinical trials. World J Clin Oncol 13, 388–411 (2022).

2. C. Blank, I. Brown, A. C. Peterson, M. Spiotto, Y. Iwai, T. Honjo, T. F. Gajewski, PD-L1/B7H-1 Inhibits the Effector Phase of Tumor Rejection by T Cell Receptor (TCR) Transgenic CD8+ T Cells. Cancer Research 64, 1140–1145 (2004).

3. M. J. Butte, M. E. Keir, T. B. Phamduy, G. J. Freeman, A. H. Sharpe, PD-L1 interacts specifically with B7-1 to inhibit T cell proliferation. Immunity 27, 111–122 (2007).

4. A. H. Sharpe, K. E. Pauken, The diverse functions of the PD1 inhibitory pathway. Nat Rev Immunol 18, 153–167 (2018).

5. A. Ribas, O. Hamid, A. Daud, F. S. Hodi, J. D. Wolchok, R. Kefford, A. M. Joshua, A. Patnaik, W.-J. Hwu, J. S. Weber, T. C. Gangadhar, P. Hersey, R. Dronca, R. W. Joseph, H. Zarour, B. Chmielowski, D. P. Lawrence, A. Algazi, N. A. Rizvi, B. Hoffner, C. Mateus, K. Gergich, J. A. Lindia, M. Giannotti, X. N. Li, S. Ebbinghaus, S. P. Kang, C. Robert, Association of Pembrolizumab With Tumor Response and Survival Among Patients With Advanced Melanoma. JAMA 315, 1600–1609 (2016).

6. C. Robert, J. Schachter, G. V. Long, A. Arance, J. J. Grob, L. Mortier, A. Daud, M. S. Carlino, C. McNeil, M. Lotem, J. Larkin, P. Lorigan, B. Neyns, C. U. Blank, O. Hamid, C. Mateus, R. Shapira-Frommer, M. Kosh, H. Zhou, N. Ibrahim, S. Ebbinghaus, A. Ribas, KEYNOTE-006 investigators, Pembrolizumab versus Ipilimumab in Advanced Melanoma. N Engl J Med 372, 2521–2532 (2015).

7. H. Borghaei, L. Paz-Ares, L. Horn, D. R. Spigel, M. Steins, N. E. Ready, L. Q. Chow, E. E. Vokes, E. Felip, E. Holgado, F. Barlesi, M. Kohlhäufl, O. Arrieta, M. A. Burgio, J. Fayette, H. Lena, E. Poddubskaya, D. E. Gerber, S. N. Gettinger, C. M. Rudin, N. Rizvi, L. Crinò, G. R. Blumenschein, S. J. Antonia, C. Dorange, C. T. Harbison, F. G. Finckenstein, J. R. Brahmer, Nivolumab versus Docetaxel in Advanced Non-squamous Non-small Cell Lung Cancer. N Engl J Med 373, 1627–1639 (2015).

8. R. J. Motzer, B. Escudier, D. F. McDermott, S. George, H. J. Hammers, S. Srinivas, S. S. Tykodi, J. A. Sosman, G. Procopio, E. R. Plimack, D. Castellano, T. K. Choueiri, H. Gurney, F. Donskov, P. Bono, J. Wagstaff, T. C. Gauler, T. Ueda, Y. Tomita, F. A. Schutz, C. Kollmannsberger, J. Larkin, A. Ravaud, J. S. Simon, L.-A. Xu, I. M. Waxman, P. Sharma, Nivolumab versus Everolimus in Advanced Renal Cell Carcinoma. N Engl J Med 373, 1803–1813 (2015).

9. R. L. Ferris, G. Blumenschein, J. Fayette, J. Guigay, A. D. Colevas, L. Licitra, K. Harrington, S. Kasper, E. E. Vokes, C. Even, F. Worden, N. F. Saba, L. C. Iglesias Docampo, R. Haddad, T. Rordorf, N. Kiyota, M. Tahara, M. Monga, M. Lynch, W. J. Geese, J. Kopit, J. W. Shaw, M. L. Gillison, Nivolumab for Recurrent Squamous-Cell Carcinoma of the Head and Neck. N Engl J Med 375, 1856–1867 (2016).

10. T. Y. Seiwert, B. Burtness, R. Mehra, J. Weiss, R. Berger, J. P. Eder, K. Heath, T. McClanahan, J. Lunceford, C. Gause, J. D. Cheng, L. Q. Chow, Safety and clinical activity of pembrolizumab for treatment of recurrent or metastatic squamous cell carcinoma of the head and neck (KEYNOTE-012): an open-label, multicentre, phase 1b trial. Lancet Oncol 17, 956–965 (2016).

11. R. Mehra, T. Y. Seiwert, S. Gupta, J. Weiss, I. Gluck, J. P. Eder, B. Burtness, M. Tahara, B. Keam, H. Kang, K. Muro, R. Geva, H. C. Chung, C.-C. Lin, D. Aurora-Garg, A. Ray, K. Pathiraja, J. Cheng, L. Q. M. Chow, R. Haddad, Efficacy and safety of pembrolizumab in recurrent/metastatic head and neck squamous cell carcinoma: pooled analyses after long-term follow-up in KEYNOTE-012. Br J Cancer 119, 153–159 (2018).

12. C. Liu, W. Peng, C. Xu, Y. Lou, M. Zhang, J. A. Wargo, J. Q. Chen, H. S. Li, S. S. Watowich, Y. Yang, D. T. Frederick, Z. A. Cooper, R. M. Mbofung, M. Whittington, K. T. Flaherty, S. E. Woodman, M. A. Davies, L. G. Radvanyi, W. W. Overwijk, G. Lizée, P. Hwu, BRAF Inhibition Increases Tumor Infiltration by T cells and Enhances the Anti-tumor Activity of Adoptive Immunotherapy in Mice. Clin Cancer Res 19, 393–403 (2013).

13. P. Sharma, S. Hu-Lieskovan, J. A. Wargo, A. Ribas, Primary, Adaptive and Acquired Resistance to Cancer Immunotherapy. Cell 168, 707–723 (2017).

14. A. C. Huang, M. A. Postow, R. J. Orlowski, R. Mick, B. Bengsch, S. Manne, W. Xu, S. Harmon, J. R. Giles, B. Wenz, M. Adamow, D. Kuk, K. S. Panageas, C. Carrera, P. Wong, F. Quagliarello, B. Wubbenhorst, K. D’Andrea, K. E. Pauken, R. S. Herati, R. P. Staupe, J. M. Schenkel, S. McGettigan, S. Kothari, S. M. George, R. H. Vonderheide, R. K. Amaravadi, G. C. Karakousis, L. M. Schuchter, X. Xu, K. L. Nathanson, J. D. Wolchok, T. C. Gangadhar, E. J. Wherry, T-cell invigoration to tumour burden ratio associated with anti-PD-1 response. Nature 545, 60–65 (2017).

15. L. Martínez-Lostao, A. Anel, J. Pardo, How Do Cytotoxic Lymphocytes Kill Cancer Cells? Clin Cancer Res 21, 5047–5056 (2015).

16. A. S. Rathore, S. Kumar, R. Konwar, A. Makker, M. P. S. Negi, M. M. Goel, CD3+, CD4+ & CD8+ tumour infiltrating lymphocytes (TILs) are predictors of favourable survival outcome in infiltrating ductal carcinoma of breast. Indian J Med Res 140, 361–369 (2014).

17. Y. K. So, S.-J. Byeon, B. M. Ku, Y. H. Ko, M.-J. Ahn, Y.-I. Son, M. K. Chung, An increase of CD8+ T cell infiltration following recurrence is a good prognosticator in HNSCC. Scientific Reports 10, 20059 (2020).

18. Y. Zhao, D. Chen, J. Yin, J. Xie, C.-Y. Sun, M. Lu, Comprehensive Analysis of Tumor Immune Microenvironment Characteristics for the Prognostic Prediction and Immunotherapy of Oral Squamous Cell Carcinoma. Front Genet 13, 788580 (2022).

19. D. B. Flies, S. Langermann, C. Jensen, M. A. Karsdal, N. Willumsen, Regulation of tumor immunity and immunotherapy by the tumor collagen extracellular matrix. Front Immunol 14, 1199513 (2023).

20. T. Courau, D. Nehar-Belaid, L. Florez, B. Levacher, T. Vazquez, F. Brimaud, B. Bellier, D. Klatzmann, TGF-β and VEGF cooperatively control the immunotolerant tumor environment and the efficacy of cancer immunotherapies. JCI Insight 1, e85974 (2016).

21. O. E. Rahma, F. S. Hodi, The Intersection between Tumor Angiogenesis and Immune Suppression. Clin Cancer Res 25, 5449–5457 (2019).

22. C. M. Bucks, J. A. Norton, A. C. Boesteanu, Y. M. Mueller, P. D. Katsikis, Chronic Antigen Stimulation Alone Is Sufficient to Drive CD8+ T Cell Exhaustion1. The Journal of Immunology 182, 6697–6708 (2009).

23. Y. Jiang, Y. Li, B. Zhu, T-cell exhaustion in the tumor microenvironment. Cell Death Dis 6, e1792–e1792 (2015).

24. X. Li, Y. Xiang, F. Li, C. Yin, B. Li, X. Ke, WNT/β-Catenin Signaling Pathway Regulating T Cell-Inflammation in the Tumor Microenvironment. Front Immunol 10, 2293 (2019).

25. M. J. Pittet, M. D. Pilato, C. Garris, T. R. Mempel, Dendritic cells as shepherds of T cell immunity in cancer. Immunity 56, 2218–2230 (2023).

26. M. Mikucki, D. Fisher, J. Matsuzaki, J. Skitzki, N. Gaulin, J. Muhitch, A. Ku, J. Frelinger, K. Odunsi, T. Gajewski, A. Luster, S. Evans, Non-redundant Requirement for CXCR3 Signaling during Tumoricidal T Cell Trafficking across Tumor Vascular Checkpoints. Nat Commun 6, 7458 (2015).

27. M. T. Chow, A. J. Ozga, R. L. Servis, D. T. Frederick, J. A. Lo, D. E. Fisher, G. J. Freeman, G. M. Boland, A. D. Luster, Intratumoral activity of the CXCR3 chemokine system is required for the efficacy of anti-PD-1 therapy. Immunity 50, 1498–1512.e5 (2019).

28. K. Kohli, V. G. Pillarisetty, T. S. Kim, Key chemokines direct migration of immune cells in solid tumors. Cancer Gene Therapy 29, 10 (2021).

29. A. J. Ozga, M. T. Chow, A. D. Luster, Chemokines and the immune response to cancer. Immunity 54, 859–874 (2021).

30. H. Bronger, J. Singer, C. Windmüller, U. Reuning, D. Zech, C. Delbridge, J. Dorn, M. Kiechle, B. Schmalfeldt, M. Schmitt, S. Avril, CXCL9 and CXCL10 predict survival and are regulated by cyclooxygenase inhibition in advanced serous ovarian cancer. Br J Cancer 115, 553–563 (2016).

31. I. G. House, P. Savas, J. Lai, A. X. Y. Chen, A. J. Oliver, Z. L. Teo, K. L. Todd, M. A. Henderson, L. Giuffrida, E. V. Petley, K. Sek, S. Mardiana, T. N. Gide, C. Quek, R. A. Scolyer, G. V. Long, J. S. Wilmott, S. Loi, P. K. Darcy, P. A. Beavis, Macrophage-Derived CXCL9 and CXCL10 Are Required for Antitumor Immune Responses Following Immune Checkpoint Blockade. Clinical Cancer Research 26, 487–504 (2020).

32. H. Prizant, N. Patil, S. Negatu, N. Bala, A. McGurk, S. A. Leddon, A. Hughson, T. D. McRae, Y.-R. Gao, A. M. Livingstone, J. R. Groom, A. D. Luster, D. J. Fowell, CXCL10+ peripheral activation niches couple preferred sites of Th1 entry with optimal APC encounter. Cell Rep 36, 109523 (2021).

33. M. Wendel, I. E. Galani, E. Suri-Payer, A. Cerwenka, Natural killer cell accumulation in tumors is dependent on IFN-gamma and CXCR3 ligands. Cancer Res 68, 8437–8445 (2008).

34. C. Yue, S. Shen, J. Deng, S. J. Priceman, W. Li, A. Huang, H. Yu, STAT3 in CD8+ T Cells Inhibits Their Tumor Accumulation by Downregulating CXCR3/CXCL10 Axis. Cancer Immunology Research 3, 864–870 (2015).

35. M. Ayers, J. Lunceford, M. Nebozhyn, E. Murphy, A. Loboda, D. R. Kaufman, A. Albright, J. D. Cheng, S. P. Kang, V. Shankaran, S. A. Piha-Paul, J. Yearley, T. Y. Seiwert, A. Ribas, T. K. McClanahan, IFN-γ–related mRNA profile predicts clinical response to PD-1 blockade. J Clin Invest 127, 2930–2940 (2017).

36. I. G. House, P. Savas, J. Lai, A. X. Y. Chen, A. J. Oliver, Z. L. Teo, K. L. Todd, M. A. Henderson, L. Giuffrida, E. V. Petley, K. Sek, S. Mardiana, T. N. Gide, C. Quek, R. A. Scolyer, G. V. Long, J. S. Wilmott, S. Loi, P. K. Darcy, P. A. Beavis, Macrophage-Derived CXCL9 and CXCL10 Are Required for Antitumor Immune Responses Following Immune Checkpoint Blockade. Clin Cancer Res 26, 487–504 (2020).

37. R. Reschke, J. Yu, B. A. Flood, E. F. Higgs, K. Hatogai, T. F. Gajewski, Immune cell and tumor cell-derived CXCL10 is indicative of immunotherapy response in metastatic melanoma. J Immunother Cancer 9, e003521 (2021).

38. Y. Zohar, G. Wildbaum, R. Novak, A. L. Salzman, M. Thelen, R. Alon, Y. Barsheshet, C. L. Karp, N. Karin, CXCL11-dependent induction of FOXP3-negative regulatory T cells suppresses autoimmune encephalomyelitis. J Clin Invest 124, 2009–2022 (2014).

39. D. J. Bangs, A. Tsitsiklis, Z. Steier, S. W. Chan, J. Kaminski, A. Streets, N. Yosef, E. A. Robey, CXCR3 regulates stem and proliferative CD8+ T cells during chronic infection by promoting interactions with DCs in splenic bridging channels. Cell Reports 38, 110266 (2022).

40. M. Dong, L. Lu, H. Xu, Z. Ruan, DC-derived CXCL10 promotes CTL activation to suppress ovarian cancer. Transl Res 272, 126–139 (2024).

41. H. Sung, J. Ferlay, R. L. Siegel, M. Laversanne, I. Soerjomataram, A. Jemal, F. Bray, Global Cancer Statistics 2020: GLOBOCAN Estimates of Incidence and Mortality Worldwide for 36 Cancers in 185 Countries. CA Cancer J Clin 71, 209–249 (2021).

42. A. R. Jethwa, S. S. Khariwala, Tobacco-related carcinogenesis in head and neck cancer. Cancer Metastasis Rev 36, 411–423 (2017).

43. A. Barsouk, J. S. Aluru, P. Rawla, K. Saginala, A. Barsouk, Epidemiology, Risk Factors, and Prevention of Head and Neck Squamous Cell Carcinoma. Med Sci (Basel) 11, 42 (2023).

44. G. D’Souza, A. R. Kreimer, R. Viscidi, M. Pawlita, C. Fakhry, W. M. Koch, W. H. Westra, M L. Gillison, Case–Control Study of Human Papillomavirus and Oropharyngeal Cancer The New England Journal of Medicine (2007), doi:10.1056/NEJMoa065497.

45. M. Lechner, J. Liu, L. Masterson, T. R. Fenton, HPV-associated oropharyngeal cancer: epidemiology, molecular biology and clinical management. Nat Rev Clin Oncol 19, 306–327 (2022).

46. I. Šimić, K. Božinović, N. Milutin Gašperov, M. Kordić, E. Pešut, L. Manojlović, M. Grce, E. Dediol, I. Sabol, Head and Neck Cancer Patients’ Survival According to HPV Status, miRNA Profiling, and Tumour Features—A Cohort Study. Int J Mol Sci 24, 3344 (2023).

47. W. N. William, X. Zhao, J. J. Bianchi, H. Y. Lin, P. Cheng, J. J. Lee, H. Carter, L. B. Alexandrov, J. P. Abraham, D. B. Spetzler, S. M. Dubinett, D. W. Cleveland, W. Cavenee, T. Davoli, S. M. Lippman, Immune evasion in HPV− head and neck precancer–cancer transition is driven by an aneuploid switch involving chromosome 9p loss. Proc Natl Acad Sci U S A 118, e2022655118 (2021).

48. X. Zhao, E. E. W. Cohen, W. N. William, J. J. Bianchi, J. P. Abraham, D. Magee, D. B. Spetzler, J. S. Gutkind, L. B. Alexandrov, W. K. Cavenee, S. M. Lippman, T. Davoli, Somatic 9p24.1 alterations in HPV– head and neck squamous cancer dictate immune microenvironment and anti-PD-1 checkpoint inhibitor activity. Proc Natl Acad Sci U S A 119, e2213835119 (2022).

49. Z. Wang, V. H. Wu, M. M. Allevato, M. Gilardi, Y. He, J. Luis Callejas-Valera, L. Vitale-Cross, D. Martin, P. Amornphimoltham, J. Mcdermott, B. S. Yung, Y. Goto, A. A. Molinolo, A. B. Sharabi, E. E. W. Cohen, Q. Chen, J. G. Lyons, L. B. Alexandrov, J. S. Gutkind, Syngeneic animal models of tobacco-associated oral cancer reveal the activity of in situ anti-CTLA-4. Nature Communications 10, 5546 (2019).

50. N. P. Judd, C. T. Allen, A. E. Winkler, R. Uppaluri, Comparative Analysis of Tumor-Infiltrating Lymphocytes in a Syngeneic Mouse Model of Oral Cancer. Otolaryngol Head Neck Surg 147, 493–500 (2012).

51. E. Cerami, J. Gao, U. Dogrusoz, B. E. Gross, S. O. Sumer, B. A. Aksoy, A. Jacobsen, C. J. Byrne, M. L. Heuer, E. Larsson, Y. Antipin, B. Reva, A. P. Goldberg, C. Sander, N. Schultz, The cBio cancer genomics portal: an open platform for exploring multidimensional cancer genomics data. Cancer Discov 2, 401–404 (2012).

52. J. Gao, B. A. Aksoy, U. Dogrusoz, G. Dresdner, B. Gross, S. O. Sumer, Y. Sun, A. Jacobsen, R. Sinha, E. Larsson, E. Cerami, C. Sander, N. Schultz, Integrative Analysis of Complex Cancer Genomics and Clinical Profiles Using the cBioPortal. Sci Signal 6, pl1 (2013).

53. Y. Li, T. Wu, S. Gong, H. Zhou, L. Yu, M. Liang, R. Shi, Z. Wu, J. Zhang, S. Li, Analysis of the Prognosis and Therapeutic Value of the CXC Chemokine Family in Head and Neck Squamous Cell Carcinoma. Front Oncol 10, 570736 (2020).

54. Y.-C. Huang, J.-L. Huang, L.-C. Tseng, P.-H. Yu, S.-Y. Chen, C.-S. Lin, High Expression of Interferon Pathway Genes CXCL10 and STAT2 Is Associated with Activated T-Cell Signature and Better Outcome of Oral Cancer Patients. J Pers Med 12, 140 (2022).

55. P. G. Meliante, F. Zoccali, M. de Vincentiis, M. Ralli, C. Petrella, M. Fiore, A. Minni, C. Barbato, Diagnostic Predictors of Immunotherapy Response in Head and Neck Squamous Cell Carcinoma. Diagnostics 13 (2023), doi:10.3390/diagnostics13050862.

56. R. S. Herbst, J.-C. Soria, M. Kowanetz, G. D. Fine, O. Hamid, M. S. Gordon, J. A. Sosman, D. F. McDermott, J. D. Powderly, S. N. Gettinger, H. E. K. Kohrt, L. Horn, D. P. Lawrence, S. Rost, M. Leabman, Y. Xiao, A. Mokatrin, H. Koeppen, P. S. Hegde, I. Mellman, D. S. Chen, F. S. Hodi, Predictive correlates of response to the anti-PD-L1 antibody MPDL3280A in cancer patients. Nature 515, 563–567 (2014).

57. J. E. Rosenberg, J. Hoffman-Censits, T. Powles, M. S. van der Heijden, A. V. Balar, A. Necchi, N. Dawson, P. H. O’Donnell, A. Balmanoukian, Y. Loriot, S. Srinivas, M. M. Retz, P. Grivas, R. W. Joseph, M. D. Galsky, M. T. Fleming, D. P. Petrylak, J. L. Perez-Gracia, H. A. Burris, D. Castellano, C. Canil, J. Bellmunt, D. Bajorin, D. Nickles, R. Bourgon, G. M. Frampton, N. Cui, S. Mariathasan, O. Abidoye, G. D. Fine, R. Dreicer, Atezolizumab in patients with locally advanced and metastatic urothelial carcinoma who have progressed following treatment with platinum-based chemotherapy: a single-arm, multicentre, phase 2 trial. Lancet 387, 1909–1920 (2016).

58. N. Au-Yeung, R. Mandhana, C. M. Horvath, Transcriptional regulation by STAT1 and STAT2 in the interferon JAK-STAT pathway. JAKSTAT 2, e23931 (2013).

59. B. K. Roberts, G. Collado, B. J. Barnes, Role of interferon regulatory factor 5 (IRF5) in tumor progression: Prognostic and therapeutic potential. Biochim Biophys Acta Rev Cancer 1879, 189061 (2024).

60. K. Tretina, E.-S. Park, A. Maminska, J. D. MacMicking, Interferon-induced guanylate-binding proteins: Guardians of host defense in health and disease. J Exp Med 216, 482–500 (2019).

61. Z. Tu, K. Li, Q. Ji, Y. Huang, S. Lv, J. Li, L. Wu, K. Huang, X. Zhu, Pan-cancer analysis: predictive role of TAP1 in cancer prognosis and response to immunotherapy. BMC Cancer 23, 133 (2023).

62. W. M. Lydiatt, S. G. Patel, B. O’Sullivan, M. S. Brandwein, J. A. Ridge, J. C. Migliacci, A. M. Loomis, J. P. Shah, Head and Neck cancers-major changes in the American Joint Committee on cancer eighth edition cancer staging manual. CA Cancer J Clin 67, 122–137 (2017).

63. D. Aran, Z. Hu, A. J. Butte, xCell: digitally portraying the tissue cellular heterogeneity landscape. Genome Biol 18, 220 (2017).

64. J. T. Chang, E. J. Wherry, A. W. Goldrath, Molecular regulation of effector and memory T cell differentiation. Nat Immunol 15, 1104–1115 (2014).

65. J. C. Sun, J. N. Beilke, L. L. Lanier, Adaptive immune features of natural killer cells. Nature 457, 557–561 (2009).

66. M. Nouri-Shirazi, J. Banchereau, D. Bell, S. Burkeholder, E. T. Kraus, J. Davoust, K. A. Palucka, Dendritic cells capture killed tumor cells and present their antigens to elicit tumor-specific immune responses. J Immunol 165, 3797–3803 (2000).

67. R. Saddawi-Konefka, A. O’Farrell, F. Faraji, L. Clubb, M. M. Allevato, S. M. Jensen, B. S. Yung, Z. Wang, V. H. Wu, N.-A. Anang, R. A. Msari, S. Schokrpur, I. F. Pietryga, A. A. Molinolo, J. P. Mesirov, A. B. Simon, B. A. Fox, J. D. Bui, A. Sharabi, E. E. W. Cohen, J. A. Califano, J. S. Gutkind, Lymphatic-preserving treatment sequencing with immune checkpoint inhibition unleashes cDC1-dependent antitumor immunity in HNSCC. Nat Commun 13, 4298 (2022).

68. S. J. Allen, D. J. Hamel, T. M. Handel, A RAPID AND EFFICIENT WAY TO OBTAIN MODIFIED CHEMOKINES FOR FUNCTIONAL AND BIOPHYSICAL STUDIES. Cytokine 55, 168–173 (2011).

69. L. Zhou, Z. Zeng, A. M. Egloff, F. Zhang, F. Guo, K. M. Campbell, P. Du, J. Fu, P. Zolkind, X. Ma, Z. Zhang, Y. Zhang, X. Wang, S. Gu, R. Riley, Y. Nakahori, J. Keegan, R. Haddad, J. D. Schoenfeld, O. Griffith, R. T. Manguso, J. A. Lederer, X. S. Liu, R. Uppaluri, Checkpoint blockade-induced CD8+ T cell differentiation in head and neck cancer responders. J Immunother Cancer 10, e004034 (2022).

70. J. R. Groom, A. D. Luster, CXCR3 in T cell function. Exp Cell Res 317, 620–631 (2011).

71. J. R. Groom, J. Richmond, T. T. Murooka, E. W. Sorensen, J. H. Sung, K. Bankert, U. H. von Andrian, J. J. Moon, T. R. Mempel, A. D. Luster, CXCR3 chemokine receptor-ligand interactions in the lymph node optimize CD4+ T helper 1 cell differentiation. Immunity 37, 1091 (2012).

72. H. Huang, S. Hao, F. Li, Z. Ye, J. Yang, J. Xiang, CD4+ Th1 cells promote CD8+ Tc1 cell survival, memory response, tumor localization and therapy by targeted delivery of interleukin 2 via acquired pMHC I complexes. Immunology 120, 148–159 (2007).

73. M. J. Ekkens, D. J. Shedlock, E. Jung, A. Troy, E. L. Pearce, H. Shen, E. J. Pearce, Th1 and Th2 cells help CD8 T-cell responses. Infect Immun 75, 2291–2296 (2007).

74. J. Kim, J. S. Kim, H. K. Lee, H. S. Kim, E. J. Park, J. E. Choi, Y. J. Choi, B. R. Shin, E. Y. Kim J. T. Hong, Y. Kim, S.-B. Han, CXCR3-deficient natural killer cells fail to migrate to B16F10 melanoma cells. Int Immunopharmacol 63, 66–73 (2018).

75. G. T. Clifton, M. Rothenberg, P. A. Ascierto, G. Begley, M. Cecchini, J. P. Eder, F. Ghiringhelli, A. Italiano, M. Kochetkova, R. Li, F. Mechta-Grigoriou, S. I. Pai, P. Provenzano, E. Puré, A. Ribas, K. A. Schalper, W. H. Fridman, Developing a definition of immune exclusion in cancer: results of a modified Delphi workshop. J Immunother Cancer 11, e006773 (2023).

76. A. Tiwari, T. Oravecz, L. A. Dillon, A. Italiano, L. Audoly, W. H. Fridman, G. T. Clifton, Towards a consensus definition of immune exclusion in cancer. Front Immunol 14, 1084887 (2023).

77. J. Galon, H. K. Angell, D. Bedognetti, F. M. Marincola, The Continuum of Cancer Immunosurveillance: Prognostic, Predictive, and Mechanistic Signatures. Immunity 39, 11–26 (2013).

78. R. Galandrini, R. De Maria, M. Piccoli, L. Frati, A. Santoni, CD44 triggering enhances human NK cell cytotoxic functions. J Immunol 153, 4399–4407 (1994).

79. S. L. Sague, C. Tato, E. Puré, C. A. Hunter, The regulation and activation of CD44 by natural killer (NK) cells and its role in the production of IFN-gamma. J Interferon Cytokine Res 24, 301– 309 (2004).

80. J. Li, M. Ge, S. Lu, J. Shi, X. Li, M. Wang, J. Huang, Y. Shao, Z. Huang, J. Zhang, N. Nie, Y. Zheng, Pro-inflammatory effects of the Th1 chemokine CXCL10 in acquired aplastic anaemia. Cytokine 94, 45–51 (2017).

81. S.-Y. Yue, D. Niu, W.-M. Ma, Y. Guan, Q.-S. Liu, X.-B. Wang, Y.-Z. Xiao, J. Meng, K. Ding, L. Zhang, H.-X. Du, C.-Z. Liang, The CXCL10/CXCR3 axis regulates Th1 cell differentiation and migration in experimental autoimmune prostatitis through the PI3K/AKT pathway. Andrology 12, 1408–1418 (2024).

82. A. J. Ozga, M. T. Chow, M. E. Lopes, R. L. Servis, M. Di Pilato, P. Dehio, J. Lian, T. R. Mempel, A. D. Luster, CXCL10 chemokine regulates heterogeneity of the CD8+ T cell response and viral set point during chronic infection. Immunity 55, 82–97.e8 (2022).

83. S. F. Ziegler, F. Ramsdell, M. R. Alderson, The activation antigen CD69. Stem Cells 12, 456– 465 (1994).

84. W. Zhang, J. Sloan-Lancaster, J. Kitchen, R. P. Trible, L. E. Samelson, LAT: The ZAP-70 Tyrosine Kinase Substrate that Links T Cell Receptor to Cellular Activation. Cell 92, 83–92 (1998).

85. Q. Guo, C.-Y. Wu, N. Jiang, S. Tong, J.-H. Wan, X.-Y. Xiao, P.-Y. Mei, H.-S. Liu, S.-H. Wang, Downregulation of T-cell cytotoxic marker IL18R1 promotes cancer proliferation and migration and is associated with dismal prognosis and immunity in lung squamous cell carcinoma. Front Immunol 13, 986447 (2022).

86. D. Duquette, C. Harmon, A. Zaborowski, X. Michelet, C. O’Farrelly, D. Winter, H.-F. Koay, L. Lynch, Human Granzyme K Is a Feature of Innate T Cells in Blood, Tissues, and Tumors, Responding to Cytokines Rather than TCR Stimulation. J Immunol 211, 633–647 (2023).

87. R. Upadhyay, J. A. Boiarsky, G. Pantsulaia, J. Svensson-Arvelund, M. J. Lin, A. Wroblewska, S. Bhalla, N. Scholler, A. Bot, J. M. Rossi, N. Sadek, S. Parekh, A. Lagana, A. Baccarini, M. Merad, B. D. Brown, J. D. Brody, A Critical Role for Fas-Mediated Off-Target Tumor Killing in T- ell Immunotherapy. Cancer Discov 11, 599–613 (2021).

88. A. Lepletier, V. P. Lutzky, D. Mittal, K. Stannard, T. S. Watkins, C. N. Ratnatunga, C. Smith, H. M. McGuire, R. A. Kemp, P. Mukhopadhyay, N. Waddell, M. J. Smyth, W. C. Dougall, J. J. Miles, The immune checkpoint CD96 defines a distinct lymphocyte phenotype and is highly expressed on tumor-infiltrating T cells. Immunol Cell Biol 97, 152–164 (2019).

89. S. Feng, O. Isayev, J. Werner, A. V. Bazhin, CD96 as a Potential Immune Regulator in Cancers. Int J Mol Sci 24, 1303 (2023).

90. D. G. Tantalo, A. J. Oliver, B. von Scheidt, A. J. Harrison, S. N. Mueller, M. H. Kershaw, C. Y. Slaney, Understanding T cell phenotype for the design of effective chimeric antigen receptor T cell therapies. J Immunother Cancer 9, e002555 (2021).

91. P. Graef, V. R. Buchholz, C. Stemberger, M. Flossdorf, L. Henkel, M. Schiemann, I. Drexler, T. Höfer, S. R. Riddell, D. H. Busch, Serial Transfer of Single-Cell-Derived Immunocompetence Reveals Stemness of CD8+ Central Memory T Cells. Immunity 41, 116–126 (2014).

92. K. M. Huster, V. Busch, M. Schiemann, K. Linkemann, K. M. Kerksiek, H. Wagner, D. H. Busch, Selective expression of IL-7 receptor on memory T cells identifies early CD40L-dependent generation of distinct CD8+ memory T cell subsets. Proc Natl Acad Sci U S A 101, 5610–5615 (2004).

93. A. Dunkle, I. Dzhagalov, C. Gordy, Y.-W. He, Transfer of CD8+ T cell memory using Bcl-2 as a marker. J Immunol 190, 940–947 (2013).

94. R. Muthuswamy, A. R. McGray, S. Battaglia, W. He, A. Miliotto, C. Eppolito, J. Matsuzaki, T. Takemasa, R. Koya, T. Chodon, B. D. Lichty, P. Shrikant, K. Odunsi, CXCR6 by increasing retention of memory CD8+ T cells in the ovarian tumor microenvironment promotes immunosurveillance and control of ovarian cancer. J Immunother Cancer 9, e003329 (2021).

95. D. J. Berg, N. Davidson, R. Kühn, W. Müller, S. Menon, G. Holland, L. Thompson-Snipes, M. W. Leach, D. Rennick, Enterocolitis and colon cancer in interleukin-10-deficient mice are associated with aberrant cytokine production and CD4(+) TH1-like responses. J Clin Invest 98, 1010–1020 (1996).

96. B. Neven, E. Mamessier, J. Bruneau, S. Kaltenbach, D. Kotlarz, F. Suarez, J. Masliah-Planchon, K. Billot, D. Canioni, P. Frange, I. Radford-Weiss, V. Asnafi, D. Murugan, C. Bole, P. Nitschke, O. Goulet, J.-L. Casanova, S. Blanche, C. Picard, O. Hermine, F. Rieux-Laucat, N. Brousse, F. Davi, V. Baud, C. Klein, B. Nadel, F. Ruemmele, A. Fischer, A Mendelian predisposition to B-cell lymphoma caused by IL-10R deficiency. Blood 122, 3713–3722 (2013).

97. M. Oft, Immune regulation and cytotoxic T cell activation of IL-10 agonists – Preclinical and clinical experience. Semin Immunol 44, 101325 (2019).

98. W. Cui, Y. Liu, J. S. Weinstein, J. Craft, S. M. Kaech, An interleukin-21-interleukin-10-STAT3 pathway is critical for functional maturation of memory CD8+ T cells. Immunity 35, 792–805 (2011).

99. Z. Liu, J.-Q. Liu, F. Talebian, L.-C. Wu, S. Li, X.-F. Bai, IL-27 enhances the survival of tumor antigen-specific CD8+ T cells and programs them into IL-10-producing, memory precursor-like effector cells. Eur J Immunol 43, 468–479 (2013).

100. N. McGranahan, A. J. S. Furness, R. Rosenthal, S. Ramskov, R. Lyngaa, S. K. Saini, M. Jamal-Hanjani, G. A. Wilson, N. J. Birkbak, C. T. Hiley, T. B. K. Watkins, S. Shafi, N. Murugaesu, R. Mitter, A. U. Akarca, J. Linares, T. Marafioti, J. Y. Henry, E. M. Van Allen, D. Miao, B. Schilling, D. Schadendorf, L. A. Garraway, V. Makarov, N. A. Rizvi, A. Snyder, M. D. Hellmann, T. Merghoub, J. D. Wolchok, S. A. Shukla, C. J. Wu, K. S. Peggs, T. A. Chan, S. R. Hadrup, S. A. Quezada, C. Swanton, Clonal neoantigens elicit T cell immunoreactivity and sensitivity to immune checkpoint blockade. Science 351, 1463–1469 (2016).

101. A. Del Prete, V. Salvi, A. Soriani, M. Laffranchi, F. Sozio, D. Bosisio, S. Sozzani, Dendritic cell subsets in cancer immunity and tumor antigen sensing. Cell Mol Immunol 20, 432–447 (2023).

102. P. See, C.-A. Dutertre, J. Chen, P. Günther, N. McGovern, S. E. Irac, M. Gunawan, M. Beyer, K. Händler, K. Duan, H. R. B. Sumatoh, N. Ruffin, M. Jouve, E. Gea-Mallorquí, R. C. M. Hennekam, T. Lim, C. C. Yip, M. Wen, B. Malleret, I. Low, N. B. Shadan, C. F. S. Fen, A. Tay, J. Lum, F. Zolezzi, A. Larbi, M. Poidinger, J. K. Y. Chan, Q. Chen, L. Rénia, M. Haniffa, P. Benaroch, A. Schlitzer, J. L. Schultze, E. W. Newell, F. Ginhoux, Mapping the human DC lineage through the integration of high-dimensional techniques. Science 356, eaag3009 (2017).

103. S. Spranger, D. Dai, B. Horton, T. Gajewski, Tumor-residing Batf3 dendritic cells are required for effector T cell trafficking and adoptive T cell therapy. Cancer Cell 31, 711–723.e4 (2017).

104. C. Carenza, F. Calcaterra, F. Oriolo, C. Di Vito, M. Ubezio, M. G. Della Porta, D. Mavilio, S. Della Bella, Costimulatory Molecules and Immune Checkpoints Are Differentially Expressed on Different Subsets of Dendritic Cells. Front Immunol 10, 1325 (2019).

105. K. S. Kobayashi, P. J. van den Elsen, NLRC5: a key regulator of MHC class I-dependent immune responses. Nat Rev Immunol 12, 813–820 (2012).

106. T. Ito, R. Amakawa, M. Inaba, S. Ikehara, K. Inaba, S. Fukuhara, Differential Regulation of Human Blood Dendritic Cell Subsets by IFNs1. The Journal of Immunology 166, 2961–2969 (2001).

107. M. von Locquenghien, C. Rozalén, T. Celià-Terrassa, Interferons in cancer immunoediting: sculpting metastasis and immunotherapy response. J Clin Invest 131, e143296.

108. K. Hildner, B. T. Edelson, W. E. Purtha, M. Diamond, H. Matsushita, M. Kohyama, B. Calderon, B. Schraml, E. R. Unanue, M. S. Diamond, R. D. Schreiber, T. L. Murphy, K. M. Murphy, Batf3 Deficiency Reveals a Critical Role for CD8α+ Dendritic Cells in Cytotoxic T Cell Immunity. Science 322, 1097–1100 (2008).

109. S. W. Lukowski, I. Rødahl, S. Kelly, M. Yu, J. Gotley, C. Zhou, S. Millard, S. B. Andersen, A. N. Christ, G. Belz, I. H. Frazer, J. Chandra, Absence of Batf3 reveals a new dimension of cell state heterogeneity within conventional dendritic cells. iScience 24, 102402 (2021).

110. M. J. O’Melia, M. P. Manspeaker, S. N. Thomas, Tumor-draining lymph nodes are survival niches that support T cell priming against lymphatic transported tumor antigen and effects of immune checkpoint blockade in TNBC. Cancer Immunol Immunother 70, 2179–2195 (2021).

111. I. Delclaux, K. S. Ventre, D. Jones, A. W. Lund, The tumor-draining lymph node as a reservoir for systemic immune surveillance. Trends Cancer 10, 28–37 (2024).

112. J. M. Schenkel, R. H. Herbst, D. Canner, A. Li, M. Hillman, S.-L. Shanahan, G. Gibbons, O. C. Smith, J. Y. Kim, P. Westcott, W. L. Hwang, W. A. Freed-Pastor, G. Eng, M. S. Cuoco, P. Rogers, J. K. Park, M. L. Burger, O. Rozenblatt-Rosen, L. Cong, K. E. Pauken, A. Regev, T. Jacks, Conventional type I dendritic cells maintain a reservoir of proliferative tumor-antigen specific TCF-1+ CD8+ T cells in tumor-draining lymph nodes. Immunity 54, 2338–2353.e6 (2021).

113. K. A. Connolly, M. Kuchroo, A. Venkat, A. Khatun, J. Wang, I. William, N. I. Hornick, B. L. Fitzgerald, M. Damo, M. Y. Kasmani, C. Cui, E. Fagerberg, I. Monroy, A. Hutchins, J. F. Cheung, G. G. Foster, D. L. Mariuzza, M. Nader, H. Zhao, W. Cui, S. Krishnaswamy, N. S. Joshi, A reservoir of stem-like CD8+ T cells in the tumor-draining lymph node preserves the ongoing antitumor immune response. Science Immunology 6, eabg7836 (2021).

114. E. J. Wherry, T cell exhaustion. Nat Immunol 12, 492–499 (2011).

115. V. H. Wu, B. S. Yung, F. Faraji, R. Saddawi-Konefka, Z. Wang, A. T. Wenzel, M. J. Song, M. S. Pagadala, L. M. Clubb, J. Chiou, S. Sinha, M. Matic, F. Raimondi, T. S. Hoang, R. Berdeaux, D. A. A. Vignali, R. Iglesias-Bartolome, H. Carter, E. Ruppin, J. P. Mesirov, J. S. Gutkind, The GPCR-Gαs-PKA signaling axis promotes T cell dysfunction and cancer immunotherapy failure. Nat Immunol 24, 1318–1330 (2023).

116. C. C. Yates-Binder, M. Rodgers, J. Jaynes, A. Wells, R. J. Bodnar, T. Turner, An IP-10 (CXCL10)-derived peptide inhibits angiogenesis. PLoS One 7, e40812 (2012).

117. M. Geindreau, M. Bruchard, F. Vegran, Role of Cytokines and Chemokines in Angiogenesis in a Tumor Context. Cancers (Basel) 14, 2446 (2022).

118. N. Gao, X. Liu, J. Wu, J. Li, C. Dong, X. Wu, X. Xiao, F.-S. X. Yu, CXCL10 suppression of hem- and lymph-angiogenesis in inflamed corneas through MMP13. Angiogenesis 20, 505–518 (2017).

119. F. Kuhnert, J. R. Kirshner, G. Thurston, Dll4-Notch signaling as a therapeutic target in tumor angiogenesis. Vasc Cell 3, 20 (2011).

120. S. A. Fry, C. E. Robertson, R. Swann, M. V. Dwek, Cadherin-5: a biomarker for metastatic breast cancer with optimum efficacy in oestrogen receptor-positive breast cancers with vascular invasion. Br J Cancer 114, 1019–1026 (2016).

121. M. Inokuchi, K. Higuchi, Y. Takagi, T. Tanioka, M. Nakagawa, K. Gokita, K. Okuno, K. Kojima, Cadherin 5 Is a Significant Risk Factor for Hematogenous Recurrence and a Prognostic Factor in Locally Advanced Gastric Cancer. Anticancer Res 37, 6807–6813 (2017).

122. X. Wang, X. Zhang, C. Zhang, L. Qi, J. Liu, Plasma von Willebrand factor levels in patients with cancer: A meta-analysis. Oncol Lett 28, 399 (2024).

123. Z. Wang, Y. Goto, M. M. Allevato, V. H. Wu, R. Saddawi-Konefka, M. Gilardi, D. Alvarado, B. S. Yung, A. O’Farrell, A. A. Molinolo, U. Duvvuri, J. R. Grandis, J. A. Califano, E. E. W. Cohen, J. S. Gutkind, Disruption of the HER3-PI3K-mTOR oncogenic signaling axis and PD-1 blockade as a multimodal precision immunotherapy in head and neck cancer. Nat Commun 12, 2383 (2021).

124. P. E. Clavijo, E. C. Moore, J. Chen, R. J. Davis, J. Friedman, Y. Kim, C. Van Waes, Z. Chen, C. T. Allen, Resistance to CTLA-4 checkpoint inhibition reversed through selective elimination of granulocytic myeloid cells. Oncotarget 8, 55804–55820 (2017).

125. D. Damotte, S. Warren, J. Arrondeau, P. Boudou-Rouquette, A. Mansuet-Lupo, J. Biton, H. Ouakrim, M. Alifano, C. Gervais, A. Bellesoeur, N. Kramkimel, C. Tlemsani, B. Burroni, A. Duche, F. Letourneur, H. Si, R. Halpin, T. Creasy, R. Herbst, X. Ren, P. Morel, A. Cesano, F. Goldwasser, K. Leroy, The tumor inflammation signature (TIS) is associated with anti-PD-1 treatment benefit in the CERTIM pan-cancer cohort. J Transl Med 17, 357 (2019).

126. A. Saudemont, N. Jouy, D. Hetuin, B. Quesnel, NK cells that are activated by CXCL10 can kill dormant tumor cells that resist CTL-mediated lysis and can express B7-H1 that stimulates T cells. Blood 105, 2428–2435 (2005).

127. H. Hu, Y. Chen, S. Tan, S. Wu, Y. Huang, S. Fu, F. Luo, J. He, The Research Progress of Antiangiogenic Therapy, Immune Therapy and Tumor Microenvironment. Front Immunol 13, 802846 (2022).

128. F. Martins, L. Sofiya, G. P. Sykiotis, F. Lamine, M. Maillard, M. Fraga, K. Shabafrouz, C. Ribi, A. Cairoli, Y. Guex-Crosier, T. Kuntzer, O. Michielin, S. Peters, G. Coukos, F. Spertini, J. A. Thompson, M. Obeid, Adverse effects of immune-checkpoint inhibitors: epidemiology, management and surveillance. Nat Rev Clin Oncol 16, 563–580 (2019).

129. Q. Yin, L. Wu, L. Han, X. Zheng, R. Tong, L. Li, L. Bai, Y. Bian, Immune-related adverse events of immune checkpoint inhibitors: a review. Front Immunol 14, 1167975 (2023).

130. E. Sebestyén, N. Major, L. Bodoki, A. Makai, I. Balogh, G. Tóth, Z. Orosz, P. Árkosy, A. Vaskó, K. Hodosi, Z. Szekanecz, É. Szekanecz, Immune-related adverse events of anti-PD-1 immune checkpoint inhibitors: a single center experience. Front Oncol 13, 1252215 (2023).

131. R. J. Lim, R. Salehi-Rad, L. M. Tran, M. S. Oh, C. Dumitras, W. P. Crosson, R. Li, T. S. Patel, S. Man, C. E. Yean, J. Abascal, Z. Huang, S. L. Ong, K. Krysan, S. M. Dubinett, B. Liu, CXCL9/10- engineered dendritic cells promote T cell activation and enhance immune checkpoint blockade for lung cancer. Cell Rep Med 5, 101479 (2024).

132. B. Kong, Y. Kim, E. H. Kim, J. S. Suk, Y. Yang, mRNA: A promising platform for cancer immunotherapy. Adv Drug Deliv Rev 199, 114993 (2023).

133. G. Xu, K. Liu, X. Chen, Y. Lin, C. Yu, X. Nie, W. He, N. Karin, Y. Luan, Hydrogel-mediated tumor T cell infiltration and immune evasion to reinforce cancer immunotherapy. Nanoscale Horiz 9, 295–304 (2024).

134. T. Ngo, B. S. Stephens, M. Gustavsson, L. G. Holden, R. Abagyan, T. M. Handel, I. Kufareva, Crosslinking-guided geometry of a complete CXC receptor-chemokine complex and the basis of chemokine subfamily selectivity. PLoS Biol 18, e3000656 (2020).

135. C. T. Schafer, Q. Chen, J. J. G. Tesmer, T. M. Handel, Atypical Chemokine Receptor 3 “Senses” CXC Chemokine Receptor 4 Activation Through GPCR Kinase Phosphorylation. Mol Pharmacol 104, 174–186 (2023).

136. C. Hennig, fpc: Flexible Procedures for Clustering (2024) (available at https://cran.r-project.org/web/packages/fpc/index.html).

137. A. L. Mitchell, T. K. Attwood, P. C. Babbitt, M. Blum, P. Bork, A. Bridge, S. D. Brown, H.-Y. Chang, S. El-Gebali, M. I. Fraser, J. Gough, D. R. Haft, H. Huang, I. Letunic, R. Lopez, A. Luciani, F. Madeira, A. Marchler-Bauer, H. Mi, D. A. Natale, M. Necci, G. Nuka, C. Orengo, A. P. Pandurangan, T. Paysan-Lafosse, S. Pesseat, S. C. Potter, M. A. Qureshi, N. D. Rawlings, N. Redaschi, L. J. Richardson, C. Rivoire, G. A. Salazar, A. Sangrador-Vegas, C. J. A. Sigrist, I. Sillitoe, G. G. Sutton, N. Thanki, P. D. Thomas, S. C. E. Tosatto, S.-Y. Yong, R. D. Finn, InterPro in 2019: improving coverage, classification and access to protein sequence annotations. Nucleic Acids Res 47, D351–D360 (2019).

138. L. Y. Geer, A. Marchler-Bauer, R. C. Geer, L. Han, J. He, S. He, C. Liu, W. Shi, S. H. Bryant, The NCBI BioSystems database. Nucleic Acids Res 38, D492–496 (2010).

139. A. Subramanian, P. Tamayo, V. K. Mootha, S. Mukherjee, B. L. Ebert, M. A. Gillette, A. Paulovich, S. L. Pomeroy, T. R. Golub, E. S. Lander, J. P. Mesirov, Gene set enrichment analysis: a knowledge-based approach for interpreting genome-wide expression profiles. Proc Natl Acad Sci U S A 102, 15545–15550 (2005).

140. A. Liberzon, A. Subramanian, R. Pinchback, H. Thorvaldsdóttir, P. Tamayo, J. P. Mesirov, Molecular signatures database (MSigDB) 3.0. Bioinformatics 27, 1739–1740 (2011).

141. B. Jassal, L. Matthews, G. Viteri, C. Gong, P. Lorente, A. Fabregat, K. Sidiropoulos, J. Cook, M. Gillespie, R. Haw, F. Loney, B. May, M. Milacic, K. Rothfels, C. Sevilla, V. Shamovsky, S. Shorser, T. Varusai, J. Weiser, G. Wu, L. Stein, H. Hermjakob, P. D’Eustachio, The reactome pathway knowledgebase. Nucleic Acids Res 48, D498–D503 (2020).

142. D. N. Slenter, M. Kutmon, K. Hanspers, A. Riutta, J. Windsor, N. Nunes, J. Mélius, E. Cirillo, S. L. Coort, D. Digles, F. Ehrhart, P. Giesbertz, M. Kalafati, M. Martens, R. Miller, K. Nishida, L. Rieswijk, A. Waagmeester, L. M. T. Eijssen, C. T. Evelo, A. R. Pico, E. L. Willighagen, WikiPathways: a multifaceted pathway database bridging metabolomics to other omics research. Nucleic Acids Res 46, D661–D667 (2018).

