## Supplemental Figures and Legends for "Activating the CXCR3/CXCL10 pathway overrides tumor immune suppression by enhancing immune trafficking and effector cell priming in head and neck squamous cell carcinoma"

### Supplemental Figure Legends

**Figure S1. Top 100 differentially expressed genes between CXCL10$^{high}$ and CXCL10$^{low}$ HNSCC tumors.** Differential expression of genes between CXCL10$^{high}$ vs CXCL10$^{low}$ cohorts using Wilcoxon Rank Sum test. $Log_2$ fold-change and P value (adjusted) cut-offs are 1 and 0.05, respectively. Transcriptomic data was obtained and analyzed via cBioPortal (PanCancer Atlas, HNSCC cohort). Expression was based on mRNA levels (whole tumor). CXCR3$^{high}$/CXCL10$^{high}$ cohorts ($n$ = 128 and 132) and CXCL10$^{low}$/CXCL10$^{low}$ cohort ($n$ = 131 and 126) represent the upper 25% and lower 25% threshold for *Cxcr3* or *Cxcl10* mRNA expression respectively.

**Figure S2. Production and functional verification of CXCR3 ligands. Proportion of complete tumor clearance across independent experiments.** (**A**) Representative lane from SDS-PAGE gel demonstrating purity of ligands (hCXCL10 shown). (**B**) Representative trace with mass-to-charge ratio of purified ligand (hCXCL10) from intact protein mass spectrometry analysis. (**C**) Approximate yield per liter of biomass processed for human and murine CXCR3 ligands. (**D**) Percent of migrating hCXCR3$^+$ Jurkat cells in response to CXCR3 ligands at varying concentrations. (**E**) Bystander BRET assay utilizing βarr2-rLuc3 and membrane-bound rGFP-CAAX to measure beta-arrestin 2 (βarr2) recruitment in CXCR3$^+$ HEK293Ts after incubation in 100 nM of indicated ligand. (**F**) Proportion of mice exhibiting complete tumor clearance across multiple independent experiments. Note that there was no treatment group for anti-PD-1 alone included in experiment 20221015.

**Figure S3. Immune cells isolated from CXCL10-treated tumors exhibit increased activation.** 1.0 x $10^6$ 4MOSC1 cells were implanted into the buccal space of C57BL/6 mice on day 0. Tumors were injected on days 3, 6, and 9 with either vehicle (50 µL of PBS) or 10 µg of CXCL10 in 50 µL of PBS (per dose). (**A**) Representative immunostaining of PanCK (cyan) and NK1.1 (red) positive cells within 4MOSC1 tumors of vehicle and CXCL10-treated mice at day 10. The scale bar represents 200 µm. (**B**) Quantification of total NK1.1+ (red) cells within the tumor stroma and tumor parenchyma (panCK, cyan) ($n$ = 4 mice per group; 2 ROIs screened per tumor). (**C**) The frequency of CD44$^+$ cells amongst total CD45$^+$NK1.1$^+$ cells at day 7, 10, and 13 ($n$ = 3-4 per group; unpaired Student's $t$ test; data are represented as mean ± SEM). (**D-E**) The frequency of CD44$^+$CD62L$^-$ cells ($T_{EFF}$) (**D**) or CD44$^+$CD62L$^+$ cells ($T_{CM}$-like) (**E**) amongst total CD45$^+$CD3$^+$ CD4$^+$ cells at day 7, 10, and 13 ($n$ = 3-4 per group; unpaired Student's $t$ test; data are represented as mean ± SEM). An increase in a $T_{CM}$-like phenotype was not observed with CD4$^+$ T cells as it was with CD8$^+$ T cells. (**F**) The frequency of TIM3$^+$ cells amongst total CD45$^+$CD3$^+$ CD8$^+$ cells on day10 ($n$ = 5 per group; unpaired Student's $t$ test; data are represented as mean ± SEM). (**G**) The frequency of PD-L1$^+$ cells amongst total CD45$^+$CD11b$^+$ CD11c$^+$ cells at day 10 ($n$ = 5 per group; unpaired Student's $t$ test; data are represented as mean ± SEM). (**H**) Representative gating strategy to identify NK$^+$, CD4$^+$, and CD8$^+$ T cells associated with **Figures 3C-D**, **4B-D**, **4I-J**, and **S3C-G**.

**Figure S4. Extended list of differentially expressed genes from tumors of vehicle and CXCL10 treated mice**. 1.0 x $10^6$ 4MOSC1 cells were implanted into the buccal space of C57BL/6 mice on day 0. Tumors injected on days 3, 6, and 9 with either vehicle (50 µL of PBS) or 10 µg of CXCL10 in 50 µL of PBS. mRNA from each tumor was isolated and quantified using the NanoString nCounter PanCancer Mouse Immune Profiling gene expression platform. Rosalind software was used to analyze differentially expressed genes ($n$ = 4 per group). Differentially expressed genes in vehicle vs CXCL10-treated 4MOSC1 mice.

**Figure S5: CXCL10 treatment promotes proliferation and reduced inhibitory markers in primary OT-1 CD8+ T cells *in vitro*. Proportion of mice exhibiting complete tumor clearance amongst independent experiments.** (**A**) Schematic depicting recognition of SIINFEKL peptide by OT-1 CD8+ T cells engineered to express an ovalbumin-specific TCR. (**B**) Number of viable cells after 48 h incubation with the indicated treatment (**C**) Proportion of TIGIT$^+$, PD-1$^+$, or CTLA4$^+$ cells of the total CD45$^+$CD3$^+$CD44$^+$CD8$^+$ cells after antigen stimulation and indicated treatment *in vitro*.

| Gene | Log Ratio | p-Value | q-Value | Higher expression in |
|---|---|---|---|---|
| **CXCL11** | 6.86 | 5.51E-106 | 5.51E-102 | CXCL10HIGH |
| **CXCL9** | 5.63 | 2.98E-83 | 1.99E-79 | CXCL10HIGH |
| **GBP1** | 3.55 | 1.33E-82 | 6.68E-79 | CXCL10HIGH |
| **CXCR2P1** | 4.54 | 6.11E-78 | 2.45E-74 | CXCL10HIGH |
| **IFNG** | 3.81 | 3.63E-76 | 1.21E-72 | CXCL10HIGH |
| **IDO1** | 5.45 | 5.47E-73 | 1.57E-69 | CXCL10HIGH |
| **TAP1** | 2.32 | 1.73E-72 | 4.34E-69 | CXCL10HIGH |
| **LAP3** | 1.97 | 2.46E-71 | 5.47E-68 | CXCL10HIGH |
| **GBP4** | 3.7 | 5.18E-71 | 1.04E-67 | CXCL10HIGH |
| **IRF1** | 2.28 | 1.56E-69 | 2.85E-66 | CXCL10HIGH |
| **GBP5** | 4.55 | 3.49E-69 | 5.81E-66 | CXCL10HIGH |
| **GZMB** | 3.56 | 2.16E-66 | 3.33E-63 | CXCL10HIGH |
| **LAG3** | 3.43 | 5.60E-66 | 8.00E-63 | CXCL10HIGH |
| **STAT1** | 2.46 | 3.41E-64 | 4.55E-61 | CXCL10HIGH |
| **BATF2** | 4.05 | 1.39E-63 | 1.73E-60 | CXCL10HIGH |
| **ETV7** | 2.96 | 1.23E-61 | 1.45E-58 | CXCL10HIGH |
| **APOL3** | 2.87 | 3.18E-60 | 3.54E-57 | CXCL10HIGH |
| **CD8A** | 3.73 | 2.18E-59 | 2.30E-56 | CXCL10HIGH |
| **FASLG** | 3.4 | 3.78E-58 | 3.78E-55 | CXCL10HIGH |
| **WARS1** | 2.93 | 7.41E-58 | 7.07E-55 | CXCL10HIGH |
| **NKG7** | 3.47 | 2.55E-57 | 2.32E-54 | CXCL10HIGH |
| **IL12RB1** | 3 | 1.40E-56 | 1.22E-53 | CXCL10HIGH |
| **TRIM21** | 1.69 | 2.12E-56 | 1.77E-53 | CXCL10HIGH |
| **PSMB9** | 2.53 | 2.79E-56 | 2.24E-53 | CXCL10HIGH |
| **EPSTI1** | 3.11 | 3.23E-56 | 2.49E-53 | CXCL10HIGH |
| **IFIT3** | 3.46 | 1.65E-55 | 1.23E-52 | CXCL10HIGH |
| **SLA2** | 2.97 | 2.23E-55 | 1.59E-52 | CXCL10HIGH |
| **PML** | 1.47 | 6.38E-55 | 4.40E-52 | CXCL10HIGH |
| **CALHM6** | 3.06 | 1.13E-54 | 7.55E-52 | CXCL10HIGH |
| **APOL6** | 2.35 | 6.64E-54 | 4.29E-51 | CXCL10HIGH |
| **TRIM22** | 2.9 | 3.15E-53 | 1.97E-50 | CXCL10HIGH |
| **GZMH** | 3.73 | 7.00E-53 | 4.25E-50 | CXCL10HIGH |
| **JAKMIP1** | 2.9 | 8.64E-53 | 5.08E-50 | CXCL10HIGH |
| **PDCD1** | 2.96 | 7.88E-52 | 4.51E-49 | CXCL10HIGH |
| **GBP1P1** | 2.66 | 3.60E-51 | 2.00E-48 | CXCL10HIGH |
| **GZMA** | 3.54 | 9.25E-51 | 4.98E-48 | CXCL10HIGH |
| **UBE2L6** | 2.09 | 9.45E-51 | 4.98E-48 | CXCL10HIGH |
| **SIRPG** | 3.16 | 9.96E-51 | 5.11E-48 | CXCL10HIGH |
| **ZNF683** | 3.54 | 1.22E-50 | 6.13E-48 | CXCL10HIGH |
| **TBX21** | 2.8 | 1.52E-50 | 7.40E-48 | CXCL10HIGH |
| **CD74** | 2.72 | 3.24E-50 | 1.54E-47 | CXCL10HIGH |
| **CXCR3** | 3.22 | 4.09E-50 | 1.91E-47 | CXCL10HIGH |
| **IRF1-AS1** | 2.05 | 4.72E-50 | 2.15E-47 | CXCL10HIGH |
| **UBA7** | 2.06 | 5.30E-50 | 2.36E-47 | CXCL10HIGH |
| **KLRC1** | 2.77 | 6.15E-50 | 2.68E-47 | CXCL10HIGH |
| **PRF1** | 2.89 | 1.28E-49 | 5.45E-47 | CXCL10HIGH |
| **KLRK1** | 2.95 | 4.92E-49 | 2.05E-46 | CXCL10HIGH |
| **TNFSF13B** | 2.61 | 5.09E-49 | 2.06E-46 | CXCL10HIGH |
| **IFIH1** | 2.36 | 5.16E-49 | 2.06E-46 | CXCL10HIGH |
| **CXCR6** | 2.83 | 8.03E-49 | 3.15E-46 | CXCL10HIGH |

| Gene | Log Ratio | p-Value | q-Value | Higher expression in |
|---|---|---|---|---|
| **CIITA** | 3.04 | 1.18E-48 | 4.56E-46 | CXCL10HIGH |
| **JAK2** | 1.89 | 2.08E-48 | 7.84E-46 | CXCL10HIGH |
| **CD2** | 2.88 | 2.12E-48 | 7.85E-46 | CXCL10HIGH |
| **IFITM1** | 2.34 | 2.60E-48 | 9.47E-46 | CXCL10HIGH |
| **HLA-E** | 1.35 | 1.12E-47 | 4.00E-45 | CXCL10HIGH |
| **CCL5** | 3.11 | 1.33E-47 | 4.69E-45 | CXCL10HIGH |
| **DDX58** | 2.23 | 1.84E-47 | 6.36E-45 | CXCL10HIGH |
| **IL18BP** | 1.97 | 2.12E-47 | 7.19E-45 | CXCL10HIGH |
| **CCL4** | 2.61 | 3.13E-47 | 1.04E-44 | CXCL10HIGH |
| **SAMHD1** | 2.33 | 3.62E-47 | 1.19E-44 | CXCL10HIGH |
| **PARP14** | 1.56 | 5.23E-47 | 1.69E-44 | CXCL10HIGH |
| **STAT2** | 1.33 | 5.41E-47 | 1.72E-44 | CXCL10HIGH |
| **CD3E** | 2.68 | 8.61E-47 | 2.69E-44 | CXCL10HIGH |
| **ZBP1** | 3.37 | 1.70E-46 | 5.23E-44 | CXCL10HIGH |
| **ICOS** | 2.74 | 2.20E-46 | 6.68E-44 | CXCL10HIGH |
| **CMPK2** | 3 | 2.80E-46 | 8.36E-44 | CXCL10HIGH |
| **CD8B** | 3.45 | 2.87E-46 | 8.46E-44 | CXCL10HIGH |
| **SAMD9L** | 2.61 | 3.14E-46 | 9.11E-44 | CXCL10HIGH |
| **IFI35** | 2.04 | 7.03E-46 | 2.01E-43 | CXCL10HIGH |
| **CCR5** | 2.82 | 7.83E-46 | 2.21E-43 | CXCL10HIGH |
| **RNF213** | 1.46 | 1.16E-45 | 3.22E-43 | CXCL10HIGH |
| **B2M** | 1.64 | 3.32E-45 | 9.09E-43 | CXCL10HIGH |
| **KIR2DL4** | 2.88 | 3.87E-45 | 1.05E-42 | CXCL10HIGH |
| **HLA-DRA** | 2.63 | 4.47E-45 | 1.19E-42 | CXCL10HIGH |
| **BTN3A3** | 1.97 | 4.95E-45 | 1.30E-42 | CXCL10HIGH |
| **CD3D** | 2.79 | 5.88E-45 | 1.53E-42 | CXCL10HIGH |
| **GPR171** | 2.55 | 7.60E-45 | 1.95E-42 | CXCL10HIGH |
| **FBXO6** | 1.42 | 1.17E-44 | 2.96E-42 | CXCL10HIGH |
| **HLA-F** | 2.28 | 1.56E-44 | 3.91E-42 | CXCL10HIGH |
| **KLRC2** | 2.65 | 1.64E-44 | 4.06E-42 | CXCL10HIGH |
| **HLA-DPA1** | 2.65 | 2.17E-44 | 5.29E-42 | CXCL10HIGH |
| **GBP2** | 2.09 | 2.74E-44 | 6.60E-42 | CXCL10HIGH |
| **RSAD2** | 3.44 | 2.97E-44 | 7.08E-42 | CXCL10HIGH |
| **UBASH3A** | 2.76 | 7.19E-44 | 1.69E-41 | CXCL10HIGH |
| **HLA-B** | 2.01 | 7.64E-44 | 1.78E-41 | CXCL10HIGH |
| **TIGIT** | 2.69 | 1.14E-43 | 2.61E-41 | CXCL10HIGH |
| **XAF1** | 2.61 | 1.32E-43 | 3.00E-41 | CXCL10HIGH |
| **IL15RA** | 1.5 | 2.28E-43 | 5.12E-41 | CXCL10HIGH |
| **PSMB10** | 1.65 | 2.56E-43 | 5.70E-41 | CXCL10HIGH |
| **BTN3A1** | 1.75 | 2.94E-43 | 6.46E-41 | CXCL10HIGH |
| **NMI** | 1.22 | 2.99E-43 | 6.50E-41 | CXCL10HIGH |
| **CTLA4** | 2.61 | 4.22E-43 | 9.09E-41 | CXCL10HIGH |
| **NLRC5** | 1.75 | 6.38E-43 | 1.36E-40 | CXCL10HIGH |
| **HLA-C** | 1.76 | 8.77E-43 | 1.85E-40 | CXCL10HIGH |
| **SAMD3** | 2.52 | 1.17E-42 | 2.44E-40 | CXCL10HIGH |
| **TARP** | 2.36 | 2.72E-42 | 5.62E-40 | CXCL10HIGH |
| **PSME2** | 1.43 | 2.76E-42 | 5.63E-40 | CXCL10HIGH |
| **ISG20** | 2.02 | 4.66E-42 | 9.42E-40 | CXCL10HIGH |
| **C1QC** | 2.49 | 5.87E-42 | 1.17E-39 | CXCL10HIGH |
| **USP18** | 2.07 | 7.88E-42 | 1.56E-39 | CXCL10HIGH |

**Figure S1.**

**A**

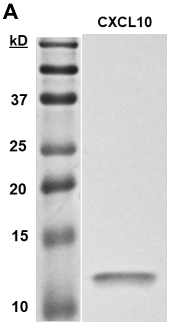


**B**

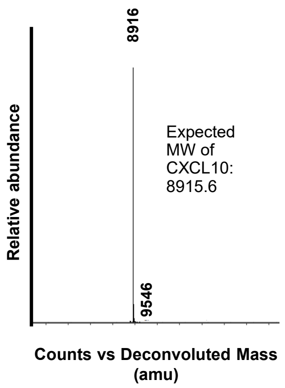


**C**

| Ligand | Yield per liter biomass (mg) |
|---|---|
| mCXCL9 | 10.5 |
| mCXCL10 | 13.5 |
| mCXCL11 | 2.1 |
| hCXCL9 | 17.2 |
| hCXCL10 | 15.0 |
| hCXCL11 | 8.5 |

**D**

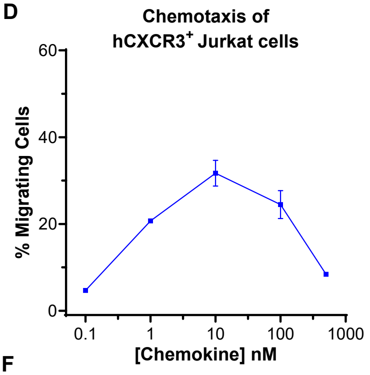


**E**

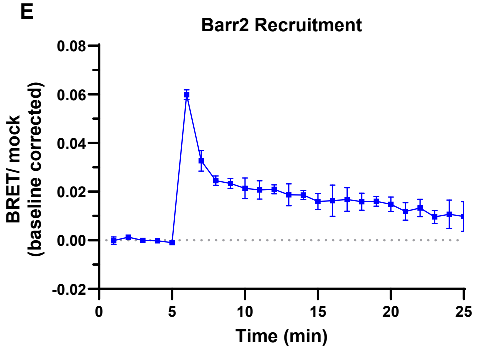


**F**

| Model | Experiment ID | Treatment Group | % Complete tumor clearance | n | Average % Complete tumor clearance |
|---|---|---|---|---|---|
| **4MOSC1** (Buccal) | 20211101* | CXCL10 | 20.00 | 5 | |
| | 20220101 | CXCL10 | 33.33 | 9 | CXCL10 24.35 |
| | 20220301 | CXCL10 | 20.00 | 5 | CXCL10 + anti-PD-1 39.17 |
| | 20220601* | CXCL10 | 20.00 | 5 | anti-PD-1 18.10 |
| | | CXCL10 + anti-PD-1 | 40.00 | 5 | |
| | | anti-PD-1 | 20.00 | 5 | |
| | 20220601 | CXCL10 | 33.33 | 6 | |
| | 20231106 | CXCL10 | 30.00 | 10 | |
| | 20220107 | CXCL10 | 30.00 | 10 | |
| | 20240715 | CXCL10 | 20.00 | 10 | |
| | | CXCL10 + anti-PD-1 | 40.00 | 10 | |
| | | anti-PD-1 | 20.00 | 10 | |
| | 20240918 | CXCL10 | 12.50 | 8 | |
| | | CXCL10 + anti-PD-1 | 37.50 | 8 | |
| | | anti-PD-1 | 14.29 | 7 | |
| **MOC1** (Flank) | 20220913* | CXCL10 (2 dose) | 10.00 | 10 | |
| | 20221015* | CXCL10 (3 dose) | 20.00 | 5 | CXCL10 (3 dose) 21.87 |
| | | CXCL10 + anti-PD-1 | 40.00 | 5 | CXCL10 + anti-PD-1 44.29 |
| | 20221130 | CXCL10 (2 dose) | 11.11 | 9 | anti-PD-1 20.54 |
| | | CXCL10 (3 dose) | 22.22 | 9 | |
| | | CXCL10 + anti-PD-1 | 50.00 | 10 | |
| | | anti-PD-1 | 12.50 | 8 | |
| | 20221130 | CXCL10 (3 dose) | 16.67 | 6 | |
| | 20230101 | CXCL10 (3 dose) | 28.57 | 7 | |
| | | CXCL10 + anti-PD-1 | 42.86 | 7 | |
| | | anti-PD-1 | 28.57 | 7 | * Pilot experiment |

**Figure S2.**

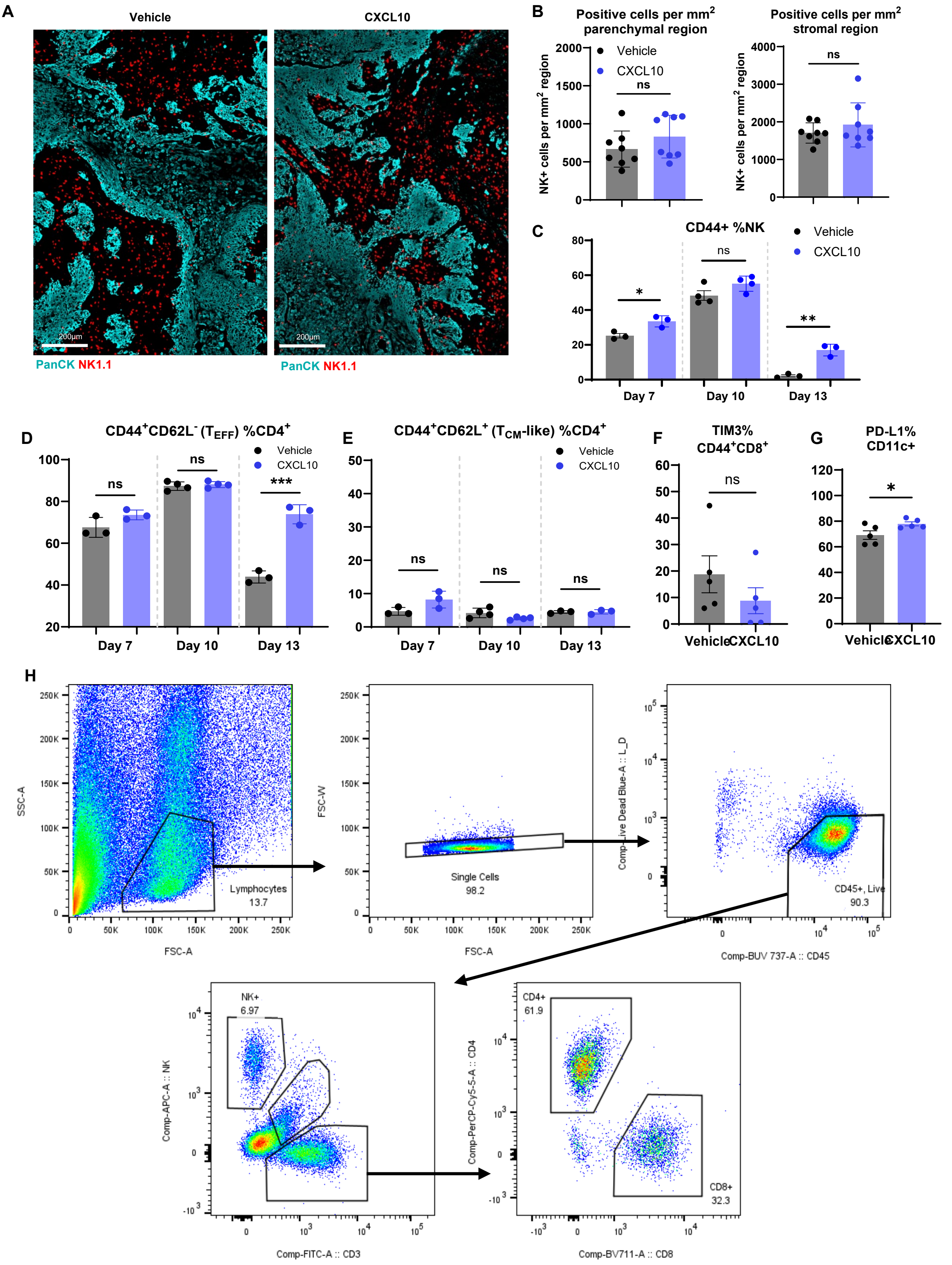

A
Vehicle
CXCL10
200µm
PanCK NK1.1
B
Positive cells per mm² parenchymal region
Positive cells per mm² stromal region
NK+ cells per mm² region
Vehicle
CXCL10
ns
C
CD44+ %NK
Day 7
Day 10
Day 13
D
CD44⁺CD62L⁻ (T_EFF) %CD4⁺
E
CD44⁺CD62L⁺ (T_CM-like) %CD4⁺
F
TIM3% CD44⁺CD8⁺
G
PD-L1% CD11c+
H
SSC-A
FSC-A
Lymphocytes 13.7
FSC-W
Single Cells 98.2
Comp-Live Dead Blue-A :: L_D
Comp-BUV 737-A :: CD45
CD45+, Live 90.3
Comp-APC-A :: NK
Comp-FITC-A :: CD3
NK+ 6.97
Comp-PerCP-Cy5-5-A :: CD4
Comp-BV711-A :: CD8
CD4+ 61.9
CD8+ 32.3


**Figure S3.**

| Name | Fold Change | p-Value |
|---|---|---|
| Aire | 3.29411 | 0.008454 |
| Amica1 | 1.40623 | 0.015678 |
| Anp32b | 1.24997 | 0.030433 |
| Anxa1 | 1.79503 | 0.031547 |
| Atf2 | 1.27996 | 0.033621 |
| Atg10 | 1.35871 | 0.033656 |
| Atg12 | 1.31462 | 0.005 |
| Atm | 1.33656 | 0.038059 |
| Batf | 1.2327 | 0.032414 |
| Bcl2 | 1.43391 | 0.003665 |
| Bid | 1.25154 | 0.034384 |
| Bst2 | 1.29917 | 0.031918 |
| C1s1 | 1.20178 | 0.012923 |
| Casp1 | 1.47831 | 0.045565 |
| Casp3 | 1.37175 | 0.011577 |
| Casp8 | 1.24014 | 0.024141 |
| Ccl2 | 1.42945 | 0.026253 |
| Ccl21a | -1.76304 | 0.020378 |
| Ccl25 | 1.82258 | 0.026259 |
| Ccr7 | 1.28389 | 0.042102 |
| Ccr8 | 1.33876 | 0.012208 |
| Cd2 | 1.26296 | 0.012408 |
| Cd207 | -3.52174 | 0.02633 |
| Cd3d | 1.35671 | 0.02064 |
| Cd3g | 1.35995 | 0.006124 |
| Cd48 | 1.2312 | 0.039091 |
| Cd59b | 1.64615 | 0.002224 |
| Cd6 | 1.50275 | 0.037716 |
| Cd69 | 1.76213 | 0.00119 |
| Cd7 | 1.22053 | 0.016641 |
| Cd86 | 1.34628 | 0.017639 |
| Cd8b1 | 2.42553 | 0.019895 |
| Cd96 | 8.67438 | 0.001574 |
| Cdh5 | -1.49532 | 0.002021 |
| Clu | -2.01175 | 0.00282 |
| Col1a1 | -1.20143 | 0.04883 |
| Col4a1 | -1.42726 | 0.002422 |
| Cspg4 | -2.04584 | 0.002603 |
| Ctla4 | 1.58195 | 0.001053 |
| Cxcl10 | 1.58422 | 0.040054 |
| Cxcl11 | 1.97931 | 0.015759 |
| Cxcl12 | -1.26809 | 0.02772 |
| Cxcl13 | 1.9781 | 0.012635 |
| Cxcl14 | -1.70374 | 0.039548 |
| Cxcl9 | 1.93139 | 0.044248 |
| Cxcr6 | 1.42559 | 0.005326 |
| Ddx60 | 1.63782 | 0.028109 |
| Dll4 | -1.62955 | 0.000528 |

| Name | Fold Change | p-Value |
|---|---|---|
| Dpp4 | 1.31961 | 0.01671 |
| Egr1 | -1.31632 | 0.020525 |
| Egr3 | -1.59155 | 0.016757 |
| Eng | -1.3452 | 0.005302 |
| Epcam | 1.27057 | 0.022537 |
| Fas | 1.23077 | 0.044811 |
| Fasl | 1.37235 | 0.000734 |
| Flt3 | 1.35877 | 0.00152 |
| Gpr183 | 1.24863 | 0.011217 |
| Gzmk | 2.87368 | 0.017236 |
| H2-M3 | 1.31429 | 0.011828 |
| H2-Ob | 1.72637 | 0.005298 |
| Hras | 1.51269 | 0.000674 |
| Icos | 1.41249 | 0.004468 |
| Ifi44 | 1.63848 | 0.023948 |
| Ifit1 | 1.91859 | 0.018442 |
| Ifit2 | 1.8737 | 0.013192 |
| Ifit3 | 1.75439 | 0.012197 |
| Ifng | 1.72581 | 0.01906 |
| Ifnl2 | 2.875 | 0.029015 |
| Igf2r | -1.27144 | 0.014245 |
| Il10 | 3.10527 | 0.021053 |
| Il12b | 1.23977 | 0.044039 |
| Il12rb1 | 1.47887 | 0.024931 |
| Il15ra | 1.55728 | 0.010566 |
| Il18r1 | 1.52986 | 4.70E-07 |
| Il18rap | 1.50572 | 0.014066 |
| Il2rb | 1.38976 | 0.035477 |
| Il7r | 1.39654 | 0.004938 |
| Irf1 | 1.21961 | 0.046856 |
| Irgm2 | 1.34728 | 0.014664 |
| Isg15 | 1.51614 | 0.010826 |
| Itga4 | 1.33736 | 0.016531 |
| Itgb1 | 1.32517 | 0.021512 |
| Kdr | -1.26115 | 0.005473 |
| Klra6 | 1.37838 | 0.039548 |
| Klrc1 | 1.47349 | 0.010835 |
| Klrc2 | 1.36667 | 0.033175 |
| Klrd1 | 1.5674 | 0.004292 |
| Klrk1 | 1.48828 | 0.009016 |
| Lbp | -1.46001 | 0.047585 |
| Lrp1 | -1.2972 | 0.009567 |
| Ly96 | 1.26472 | 0.034077 |
| Map2k1 | 1.22086 | 0.012341 |
| Map3k7 | 1.24899 | 0.037387 |
| Mfge8 | -1.34943 | 0.019646 |
| Mmp9 | -1.48419 | 0.006174 |
| Mx2 | 1.43296 | 0.017716 |
| Nfatc4 | -1.24702 | 0.01394 |
| Nlrc5 | 1.61033 | 0.003287 |

| Name | Fold Change | p-Value |
|---|---|---|
| Nlrp3 | 1.20832 | 0.032459 |
| Pdcd1 | 1.65342 | 0.03792 |
| Pdgfrb | -1.25917 | 0.024725 |
| Pou2af1 | 2.39999 | 0.044842 |
| Prkce | 1.25361 | 0.002413 |
| Psma2 | 1.24721 | 0.004736 |
| Psmb7 | 1.20604 | 0.013435 |
| Psmb8 | 1.22817 | 0.045758 |
| Psmb9 | 1.35575 | 0.014726 |
| Pvrl2 | -1.28224 | 0.023564 |
| Pycard | 1.45638 | 0.024241 |
| Ripk2 | 1.65957 | 0.003502 |
| Sell | 1.28739 | 0.041648 |
| Sh2d1a | 1.41523 | 0.000292 |
| Slamf1 | 2.2647 | 0.005106 |
| Smad2 | 1.22917 | 0.024589 |
| Tbk1 | 1.2842 | 0.016939 |
| Tek | -1.42376 | 0.014675 |
| Tgfb3 | -1.32146 | 0.00354 |
| Tie1 | -1.35427 | 0.017119 |
| Tigit | 1.37815 | 0.020684 |
| Timd4 | 1.59382 | 0.021533 |
| Tnfaip3 | 1.29121 | 0.017578 |
| Tnfrsf10b | 1.34705 | 0.018551 |
| Tnfrsf18 | 1.41059 | 0.004994 |
| Tnfsf10 | 1.45769 | 0.047425 |
| Tnfsf12 | -1.43134 | 0.019634 |
| Txk | 1.48276 | 0.025025 |
| Usp18 | 1.4497 | 0.003965 |
| Vim | -1.40089 | 0.012007 |
| Vwf | -1.62091 | 0.017623 |
| Xaf1 | 1.43289 | 0.005109 |
| Zap70 | 1.32695 | 0.017549 |
| Zbp1 | 1.32632 | 0.0165 |

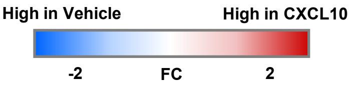


**Figure S4.**

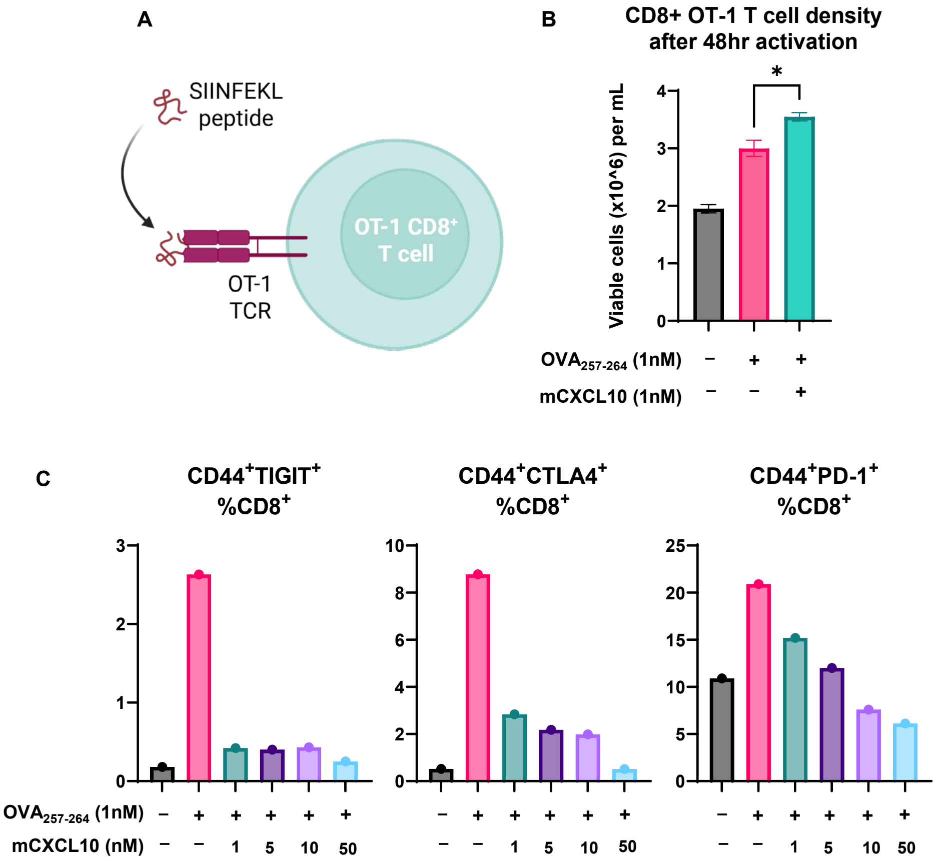
A
SIINFEKL peptide
OT-1 CD8+ T cell
OT-1 TCR
B
CD8+ OT-1 T cell density after 48hr activation
Viable cells (x10^6) per mL
OVA257-264 (1nM)
mCXCL10 (1nM)
C
CD44+TIGIT+ %CD8+
CD44+CTLA4+ %CD8+
CD44+PD-1+ %CD8+
OVA257-264 (1nM)
mCXCL10 (nM)

**Figure S5.**
